# The RNA helicase DDX1 associates with the nuclear RNA exosome and modulates R-loops

**DOI:** 10.1101/2023.04.17.537228

**Authors:** Julia L. de Amorim, Sara W. Leung, Ramona Haji-Seyed-Javadi, Yingzi Hou, David S. Yu, Homa Ghalei, Sohail Khoshnevis, Bing Yao, Anita H. Corbett

## Abstract

The RNA exosome is a ribonuclease complex that mediates both RNA processing and degradation. This complex is evolutionarily conserved, ubiquitously expressed, and required for fundamental cellular functions, including rRNA processing. The RNA exosome plays roles in regulating gene expression and protecting the genome, including modulating the accumulation of RNA-DNA hybrids (R-loops). The function of the RNA exosome is facilitated by cofactors, such as the RNA helicase MTR4, which binds/remodels RNAs. Recently, missense mutations in RNA exosome subunit genes have been linked to neurological diseases. One possibility to explain why missense mutations in genes encoding RNA exosome subunits lead to neurological diseases is that the complex may interact with cell- or tissue-specific cofactors that are impacted by these changes. To begin addressing this question, we performed immunoprecipitation of the RNA exosome subunit, EXOSC3, in a neuronal cell line (N2A) followed by proteomic analyses to identify novel interactors. We identified the putative RNA helicase, DDX1, as an interactor. DDX1 plays roles in double-strand break repair, rRNA processing, and R-loop modulation. To explore the functional connections between EXOSC3 and DDX1, we examined the interaction following double-strand breaks, and analyzed changes in R-loops in N2A cells depleted for EXOSC3 or DDX1 by DNA/RNA immunoprecipitation followed by sequencing (DRIP-Seq). We find that EXOSC3 interaction with DDX1 is decreased in the presence of DNA damage and that loss of EXOSC3 or DDX1 alters R-loops. These results suggest EXOSC3 and DDX1 interact during events of cellular homeostasis and potentially suppress unscrupulous expression of genes promoting neuronal projection.

## Introduction

The RNA exosome is a 10-subunit ribonuclease complex responsible for processing and degradation of many classes of RNA in all eukaryotes and many archaea. The ribonuclease activity of the RNA exosome is critical for both RNA quality control and precise processing of key RNAs, including ribosomal RNA (rRNA) (1–3). As illustrated in **Figure 1A**, the 10 subunits of the RNA exosome are organized into a non-catalytic cap, composed of three subunits (EXOSC1, EXOSC2, and EXOSC3), a non-catalytic core ring, comprising six subunits (EXOSC4, EXOSC5, EXOSC6, EXOSC7, EXOSC8, and EXOSC9), and one catalytic 3’-5’ exo/endoribonuclease subunit (DIS3 or DIS3L) (4–9). Most target RNAs are threaded through the cap and central channel of the RNA exosome to reach DIS3 or DIS3L for processing and/or degradation (5,10–12). Studies in yeast and other model systems have shown that the RNA exosome complex is essential (2,13–18) and ubiquitously expressed (19). Although this complex has been studied for decades, key questions, such as how the RNA exosome is targeted to specific RNAs, remain to be answered.

**Figure 1:**
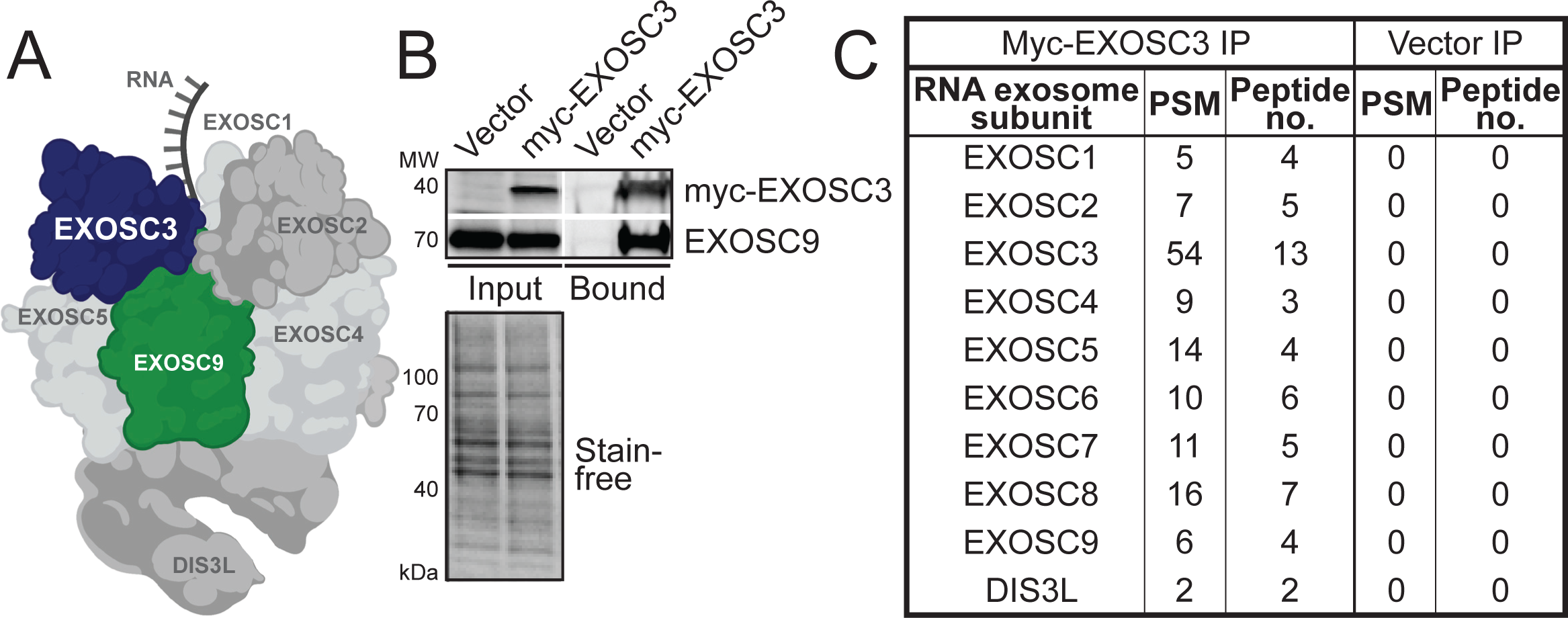
RNA exosome subunits co-immunoprecipitate with tagged EXOSC3. (A) The RNA exosome is a conserved exo/endoribonuclease complex that comprises 10 subunits. Nine of the ten subunits are structural and termed exosome components. EXOSC1, EXOSC2, and EXOSC3 make up the cap and EXOSC4, EXOSC5, EXOSC6, EXOSC7, EXOSC8, and EXOSC9 make up a barrel-shaped core. In this graphic, EXOSC3 (navy) and EXOSC9 (green) are highlighted. EXOSC6, EXOSC7, and EXOSC8 are positioned behind subunits EXOSC4, EXOSC5, and EXOSC9, and consequently are not visible. The catalytic subunit, DIS3 or DIS3L, sits at the base of the complex. This structure was created using Biorender and is based on PDB 6H25 (6). (B) EXOSC9 core subunit co-precipitates with myc-EXOSC3 from murine neuronal N2A cell line. Cells were transfected with a plasmid encoding Vector control or myc-EXOSC3 followed by immunoprecipitation using anti-myc magnetic beads. Input for Vector control and myc-EXOSC3 was probed by an anti-myc antibody, and a band corresponding to the molecular weight is detected in the Input but not Vector control for myc-EXOSC3. Input for Vector control and myc-EXOSC3 is probed by an anti-EXOSC9 antibody, and a band at the corresponding molecular weight is present in both lanes. Bound for Vector control and myc-EXOSC3 is probed by an anti-myc and an anti-EXOSC9 antibody, and a band corresponding to the molecular weight is detected in the Bound fraction for myc-EXOSC3 but not for Vector control. Stain-free blot indicates the loading of total protein in the Input. Immunoprecipitation of the myc tagged EXOSC3 copurifies with the endogenous EXOSC9 subunit. (C) Eluates of the Bound myc-EXOSC3 immunoprecipitation were analyzed by LC-MS/MS. A table shows all RNA exosome subunits detected, listing the peptide-spectrum matches (PSM) and the peptide numbers for each subunit. Vector IP serves as a control.

The RNA exosome complex processes/degrades multiple classes of RNAs in both the nucleus and the cytoplasm. The best-defined role of the RNA exosome is the processing of precursor rRNA to mature rRNA for the production of ribosomes (1,2,20–22). Other RNAs in the nucleus/nucleolus targeted by the RNA exosome include RNA from RNA/DNA hybrids, commonly known as R-loops, promoter upstream transcripts (PROMPTs), small nuclear RNAs (snRNAs), small nucleolar RNAs (snoRNAs), and other non-coding RNAs (ncRNAs), such as transcription start site (TSS)-associated antisense transcripts (xTSS-RNA) (3,23–28). In the cytoplasm, the RNA exosome targets aberrant transcripts for degradation, mRNAs for regulatory turnover, and viral RNAs as a cellular immune response (26,29–33). The RNA exosome has also been implicated in DNA double-strand break repair by homologous recombination (HR), potentially targeting aberrant transcripts produced upon DNA damage (34). The transcripts that are targeted in response to DNA damage are often within R-loop structures, which occur naturally during transcription. An accumulation of R-loops may have deleterious effects, leading to double-strand breaks and genomic instability (35). Studies suggest that the RNA exosome is poised to degrade released RNA after R-loops are unwound by RNA/DNA helicases (36).

The RNA exosome plays a critical role in cells by degrading and/or processing many transcripts in different cellular compartments. Thus, missense mutations in genes encoding structural subunits of the complex are linked to several human diseases, termed exosomopathies (37,38). The first link between the RNA exosome complex and disease described a patient with a missense mutation in *EXOSC3*, which causes the neurological disease pontocerebellar hypoplasia type 1B (PCH1B) (39). This autosomal recessive disease is characterized by severe atrophy and progressive hypoplasia of the pons and cerebellum (40–42). Since the initial report of mutations in *EXOSC3*, more mutations have been identified and described in *EXOSC1*, *EXOSC2*, *EXOSC5*, *EXOSC8*, and *EXOSC9* (16,17,43–46). All patients, with the exception of individuals with mutations in *EXOSC2*, suffer from cerebellar atrophy, at least to some extent. Patients with mutations in *EXOSC2* present with a syndromic condition that consists of short stature, hearing loss, retinitis pigmentosa, distinctive facies, and mild intellectual disability (44,47). Why mutations in genes encoding structural subunits of the RNA exosome impact the cerebellum is not at all clear.

A number of protein cofactors associate with the RNA exosome to confer RNA target specificity. Several RNA exosome cofactors that were originally identified and characterized in budding yeast are conserved in human (4,48–53). For example, the RNA exosome requires helicases for proper RNA processing as RNAs with significant secondary structure are unable to enter the central channel of the hexameric ring of the RNA exosome (5). A nuclear helicase, termed MTR4 (alternatively named SKIV2L2), interacts with the RNA exosome and facilitates processing of RNA in the nucleus (7,8). A well characterized cytoplasmic helicase, termed SKIV2L in humans or Ski2 in budding yeast, directly interacts with the RNA exosome and assists in cytoplasmic rRNA processing (49,51). The RNA exosome also interacts with nuclear scaffolding proteins, such as MPP6 (alternatively named M-phase phosphoprotein 6), and other associated ribonucleases, such as EXOSC10 (human) or Rrp6 (budding yeast) (7,8). One potential hypothesis to explain how single amino acid changes in structural subunits of the RNA exosome cause disease is that modest changes in the subunits alter interactions of the complex with cofactors required to target and subsequently process or degrade specific RNAs. As the RNA exosome is essential for fundamental processes, such as the production of mature ribosomes, a complete loss of function in patients seems unlikely. Why the majority of missense mutations identified in patients with exosomopathies cause clinical consequences most notable within regions of the brain remains unclear.

In this study, we examined the interactome of the RNA exosome in a neuronal cell line. An unbiased mass spectrometry approach identified a number of candidate binding partners. We identified the putative RNA helicase DDX1 as a protein that interacts with EXOSC3 and explored the shared functions. We found that DDX1 interacts with EXOSC3 in the nucleus, that the interaction is DNA damage-sensitive, and that the depletion of EXOSC3 or DDX1 results in significant changes in R-loops. Together, these findings suggest a novel aspect of RNA exosome function and regulation that is required for gene expression control.

## Results

### Proteomics reveal a suite of EXOSC3 interactors

To identify RNA exosome-interacting proteins in a neuronal cell line, we transiently transfected a plasmid encoding myc-tagged EXOSC3 into a murine neuroblastoma cell line (N2A) and purified co-precipitated proteins using anti-myc magnetic beads as described in Materials and Methods. Immunoblots shown in **Figure 1B** confirm that myc-EXOSC3 is enriched in the Bound fraction compared with the Vector control. The core subunit EXOSC9 of the RNA exosome co-precipitates with myc-EXOSC3, suggesting myc-tagged EXOSC3 associates with the RNA exosome complex. We then subjected the immunoprecipitates to LC-MS/MS as described in Materials and Methods. The table in **Figure 1C** shows that all subunits of the RNA exosome complex were detected in the myc-EXOSC3 immunoprecipitation as compared to samples from the control.

The RNA exosome-interacting proteins identified by mass spectrometry were analyzed using the Panther Gene Ontology (GO) program and organized by protein class (**Figure 2A**). A complete list of the interacting proteins is provided in **Table S1**. We excluded all proteins for which the peptide spectra matches (PSM) log_2_ ratio was less than or equal to zero. We examined 955 proteins for which the PSM equated to greater than zero. The largest category of the GO protein classes is “translation” containing 120 proteins and the second largest is “nucleic acid metabolism and binding” with 114 proteins. Within the latter category, all the RNA exosome subunits and some known cofactors, including MTR4 (also known as SKIV2L2) and MPP6, are present. **Figure 2B** shows cofactors (green) and potential RNA exosome-interacting candidates selected for further analysis (blue). The PSM and peptide numbers are low for even well-established cofactors, and therefore we used literary analysis and a higher PSM and peptide number to inform decisions to explore specific interactors. For this analysis, we opted to focus on Nucleic acid metabolism/binding instead of Translation because several cytoplasmic interactions between the RNA exosome and cofactors have been well characterized (49,50) and interactions between the RNA exosome and ribosome subunits have been defined (51,52).

**Figure 2:**
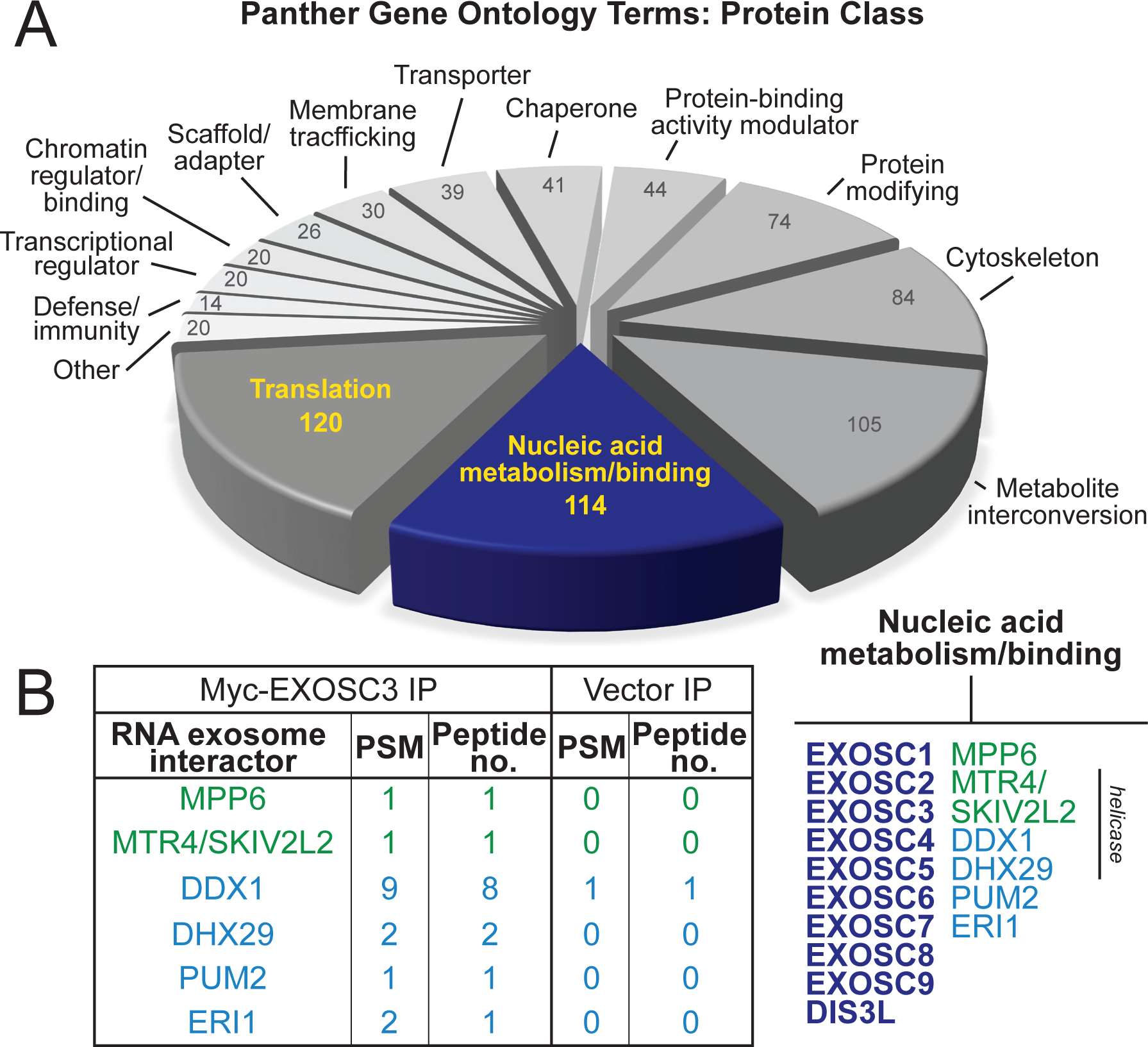
Novel EXOSC3/RNA exosome interactors identified using liquid chromatography coupled with tandem mass spectrometry (LC-MS/MS). (A) Pie chart that organizes proteins that co-precipitate with myc-EXSOSC3 from murine neuronal N2A cell line by protein class. Peptide-spectrum matches (PSM) were analyzed using a log_2_ ratio, so that any result above 0 indicates binding in myc-EXOSC3 IP over Vector IP. Results that are less than or equal to 0 were excluded from the analyses. Proteins that contained a PSM beyond the log_2_ cut-off of 0 were analyzed by Panther Gene Ontology terms. The number of proteins within a class is indicated inside the pie slices. (B) The table (left) shows selected RNA exosome interactors detected and lists the PSM and Peptide number for each protein. Vector IP serves as the control. The gene list corresponding to the proteins is provided as Supporting Information Table S1. Note that MPP6 is the protein name and MPHOSPH6 is the gene name. MTR4 is the most common name for this helicase but appears as SKIV2L2 in Table S1. A short list of Nucleic acid metabolism/binding (right) is provided. The RNA exosome subunits (bold, navy) reside in the nucleic acid metabolism/binding subcategory, together with known RNA exosome cofactors (green). Candidates investigated as novel RNA exosome interactors are listed in blue, including the putative RNA helicase, DDX1.

### Putative helicase DDX1 interacts with the RNA exosome in the nucleus

While we analyzed a number of candidates, we focused on the putative RNA helicase DDX1 for several reasons. RNA helicases play critical roles in various aspects of RNA metabolism, including RNA degradation and processing (54). Additionally, multiple conserved RNA exosome cofactors are helicases (7,51). Based on sequence homology, DDX1 is a putative helicase containing a conserved DEAD amino acid sequence motif shared by nucleic acid-unwinding, ATP-binding, DEAD-box proteins (54–56). DDX1 differs from other members of the DEAD-box family, as it includes an N-terminal SPRY protein interacting domain upstream of two helicase domains, between the phosphate-binding P-loop and the single-strand DNA binding Ia motifs (**Figure 3A**) (54). The DDX1 protein is implicated in rRNA processing (57), R-loop formation (58), and double-strand break repair (59,60).

**Figure 3:**
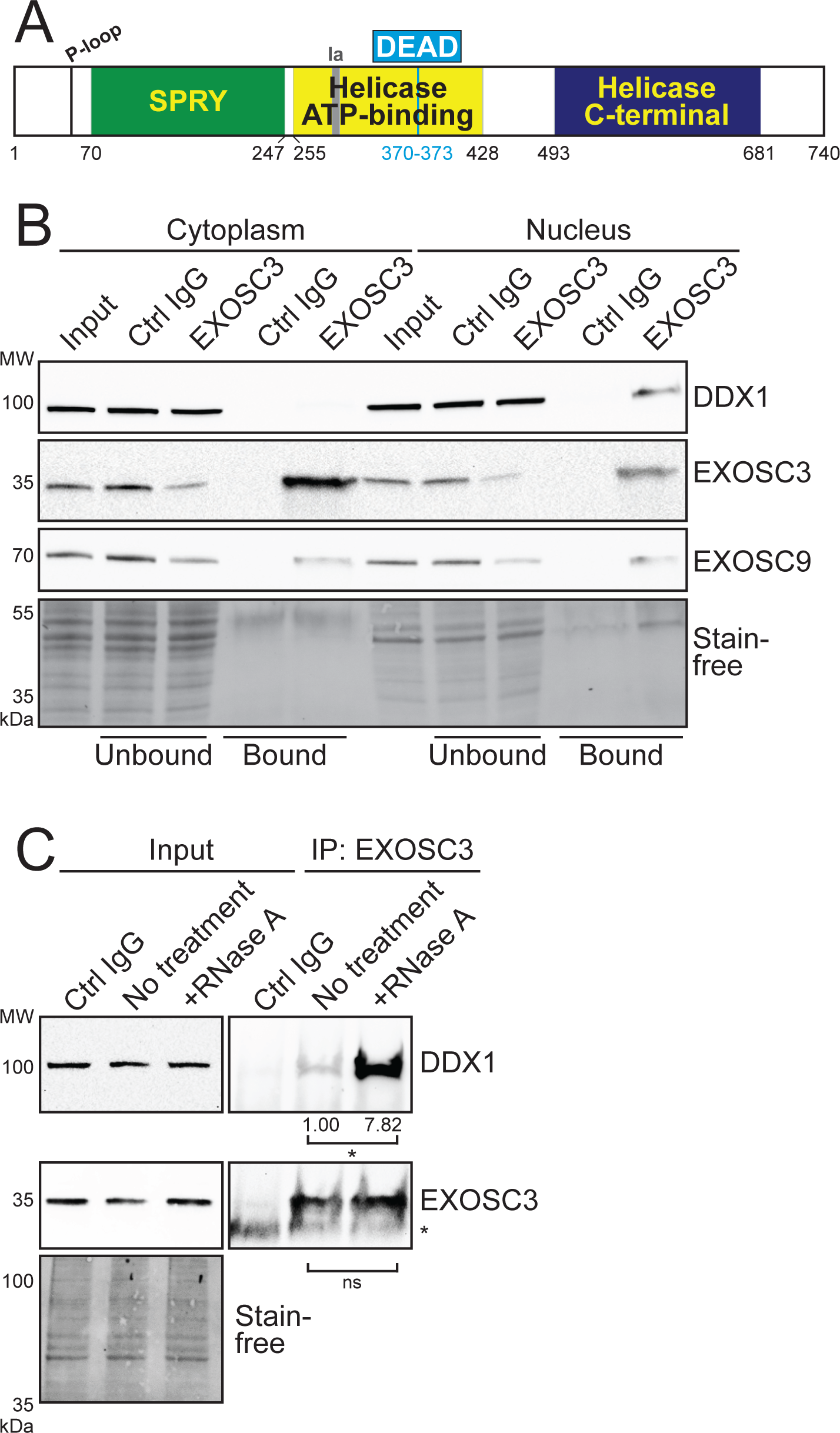
DDX1 co-immunoprecipitates with EXOSC3. (A) A graphical representation of the domain structure of human putative DEAD-box helicase DDX1. The SPRY protein interacting domain in DDX1 is located between a phosphate-binding P-loop motif and a single-strand DNA binding Ia motif, separating the motifs by 240 residues instead of the usual 20-40 residues seen in other DEAD-box proteins (54). The catalytic ATP-binding helicase and C-terminal helicase domains lie downstream of the SPRY domain. (B) DDX1 co-immunoprecipitates with EXOSC3 in the nuclear, but not cytoplasmic, fraction of N2A cell lysate. EXOSC3 was immunoprecipitated from the cytoplasmic or nuclear fraction, followed by immunoblotting using EXOSC3, EXOSC9, and DDX1 antibodies. The Input, Unbound, and Bound fractions from the EXOSC3 and control nonspecific rabbit IgG (Ctrl IgG) immunoprecipitation are shown. EXOSC9 serves as a representative of the co-purified RNA exosome subunits. Stain-free blot serves as the loading control. (C) Immunoprecipitation of EXOSC3 from the nuclear fraction was performed with RNase A treatment. EXOSC3 antibody described previously is used in the No treatment and +RNase A immunoprecipitation. Nonspecific rabbit IgG (Ctrl IgG) was used as a control. EXOSC3 and DDX1 were analyzed for the Input and bound fractions (IP: EXOSC3). The IgG light chain band is visible (asterisk) in the bound fractions probed with EXOSC3, just below the EXOSC3 band. Stain-free blot serves as a loading control for the Input. The experiment was performed in biological replicates (n=5) and bands in the bound fractions were quantified relative to the No treatment control. The values below the lanes correspond to the amount of protein quantified from the bands. The asterisk (*) below the values indicate the P-value < 0.05.

Initial studies employed epitope-tagged, transiently transfected EXOSC3. To test interactions with endogenous EXOSC3, we raised a rabbit polyclonal antibody against mouse EXOSC3. To test the specificity of this newly generated antibody, we depleted cells of EXOSC3 by transfecting N2A cells with two independent siRNA oligonucleotides that target *EXOSC3* and performed immunoblotting (**Figure S1A**). In total N2A cell lysate, the antibody detects a prominent band at the predicted size of EXOSC3 (calculated molecular weight of 29.5 kDa). This band is significantly decreased when cells are treated with either siRNA targeting *EXOSC3* as compared to the Scramble control, providing evidence for the specificity of the antibody generated.

Initial attempts to validate putative RNA exosome interacting proteins identified by mass spectrometry in whole cell lysate were unsuccessful. Thus, we considered the fact that many RNA exosome cofactors localize to specific cellular compartments (4), and we examined interactions with endogenous EXOSC3 using cellular fractionation as described in Materials and Methods. Fractionation was confirmed via immunoblotting for the cytoplasmic marker HSP90 (61) and the nuclear protein B23 (62) (**Figure S1B**). A band corresponding to the B23 is enriched in the nuclear fraction and absent in the cytoplasmic fraction, indicating efficient nuclear isolation. However, a band corresponding to HSP90 in the nuclear fraction suggests some cytoplasmic adulteration. We immunoprecipitated endogenous EXOSC3 from both the nuclear and cytoplasmic fractions, then used SDS-PAGE and immunoblotting to assess co-purification. DDX1 is detected in the Input of both the nuclear and cytoplasmic lysates, consistent with reported localization (**Figure 3B**) (63,64). However, DDX1 is present in the Bound fraction for only the Nucleus and not the Cytoplasm. DDX1 was not detected in any of the Bound fractions for Ctrl IgG samples. EXOSC9, a core component of the RNA exosome complex is detected in both the Cytoplasm and Nucleus Inputs as well as the Bound fractions, consistent with the fact that the RNA exosome complex is present in both compartments. EXOSC3 is enriched in the Bound fractions in both the Cytoplasm and Nucleus, and not in control IgG (Ctrl IgG). These data suggest a compartment-specific interaction between EXOSC3 and DDX1 in the nuclear fraction.

To assess whether the interaction detected between EXOSC3 and DDX1 is RNA-dependent, we treated the nuclear N2A cell lysate with RNase A. A urea-PAGE gel confirmed that RNase A degraded total RNA from the lysate (data not shown). We found that the treatment with RNase A increases the interaction between EXOSC3 and DDX1 with no detectable effect on the steady-state level of either protein (**Figure 3C**). This experiment suggests the interaction is not RNA-dependent and the significant increase in the interaction with RNase A treatment has been suggested to indicate a protein-protein interaction (65).

### The interaction between EXOSC3 and DDX1 decreases in response to DNA damage

The interaction between EXOSC3 and DDX1 is nuclear-specific and is increased in the absence of RNA. We sought to further characterize the interaction between the two proteins. Because both the RNA exosome and DDX1 play roles in DNA damage repair (34,59,60), we tested whether the interaction between the proteins is sensitive upon induction of DNA double-strand breaks. We treated N2A cells with 5 µM camptothecin (CPT), a topoisomerase inhibitor that induces double-strand breaks (66), or control phosphate-buffered saline (PBS) at 37°C for one hour. To confirm DNA damage, we used immunofluorescence to detect γH2AX, a classic marker of double-strand breaks (67). As shown in **Figure 4A**, after treatment with camptothecin, the γH2AX signal increases markedly compared with the PBS control. We used the EXOSC3 antibody to immunoprecipitate endogenous EXOSC3 from the nuclear fraction of cells treated with CPT and probed for DDX1 (**Figure 4B**). The interaction between EXOSC3 and DDX1 is significantly reduced following treatment with camptothecin (**Figure 4C**). In contrast, there is no change in the interaction between EXOSC3 and EXOSC9. Additionally, the compartmental localization of the interaction between EXOSC3 and DDX1 does not change upon damage (**Figure S2**). Thus, the interaction between EXOSC3 and DDX1 is substantially decreased in response to the induction of double-strand breaks.

**Figure 4:**
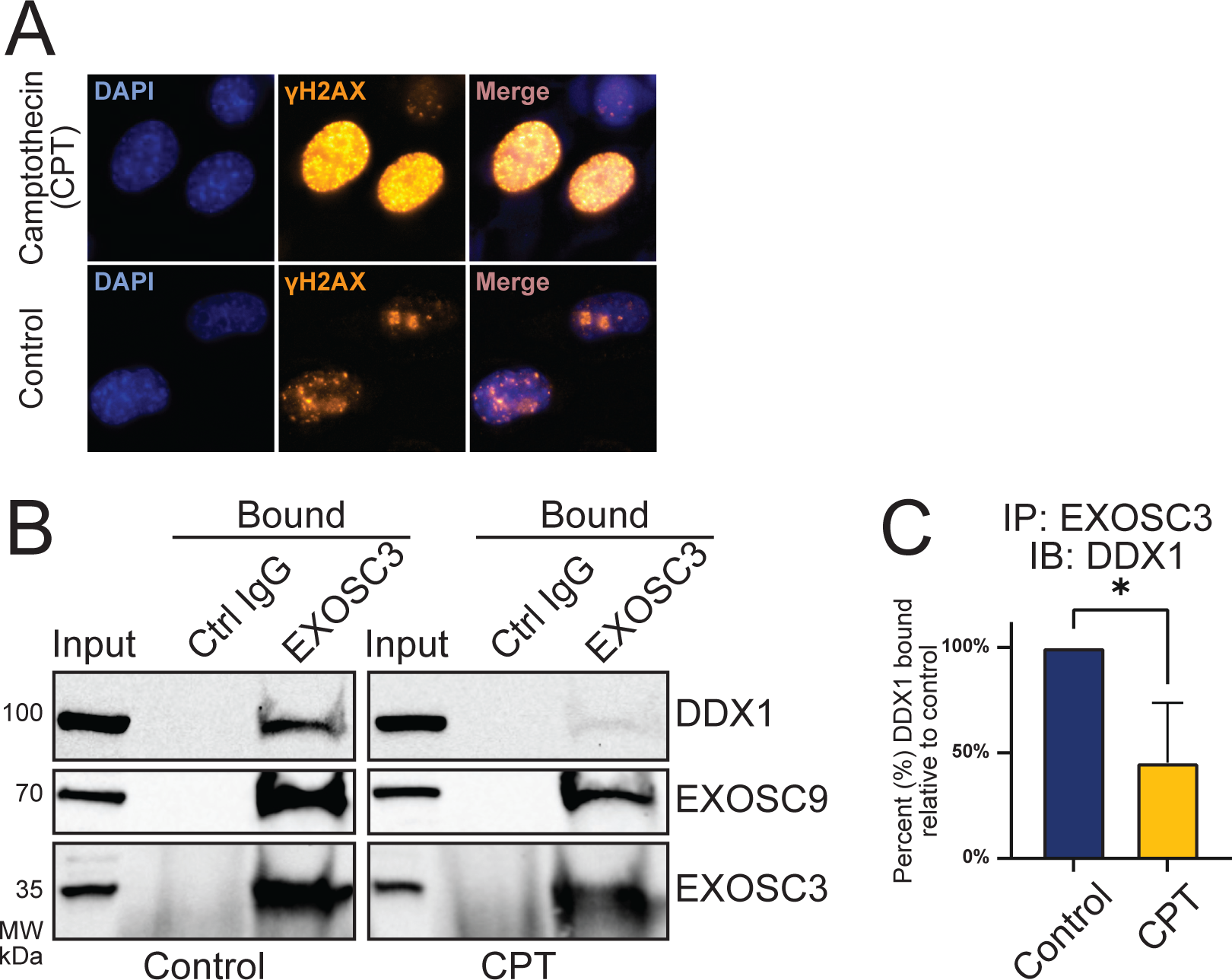
The interaction between EXOSC3 and DDX1 is sensitive to DNA damage. (A) N2A cells were treated with camptothecin (CPT) or PBS (Control), fixed, and analyzed by immunofluorescence using an antibody that detects the DNA damage marker, γH2AX. (B) The Input and immunoprecipitated samples from nuclear fractions (Bound) treated with either CPT or PBS (Control) for both EXOSC3 and control IgG (Ctrl IgG) are shown. DDX1, EXOSC9, and EXOSC3 are detected. (C) The immunoprecipitation experiment in Figure 4B was performed in biological triplicate and DDX1 bands in EXOSC3 Bound fractions were quantified. Statistical significance was calculated by a student’s T-test. Asterisk (*) represents p-value < 0.05. IP: immunoprecipitation; IB: immunoblot.

### Depletion of EXOSC3 or DDX1 results in rRNA processing defects

To explore potential shared functions of the RNA exosome and DDX1, we optimized conditions to siRNA deplete each protein from N2A cells. We transfected N2A cells with siRNA scramble control (Scramble) or EXOSC3 siRNA, then performed an immunoblot to confirm depletion (**Figure 5A**). Knockdown was quantified across the three biological replicates (**Figure 5B**). Similarly, N2A cells were transfected with DDX1 siRNA, and depletion was confirmed by immunoblot (**Figure 5C**). The knockdown was quantified across the three biological replicates (**Figure 5D**). Thus, we were able to substantially deplete each protein to less than 15% remaining (**Figure 5B, 5D**), providing a model to explore and compare the consequences of loss of each of these proteins.

**Figure 5:**
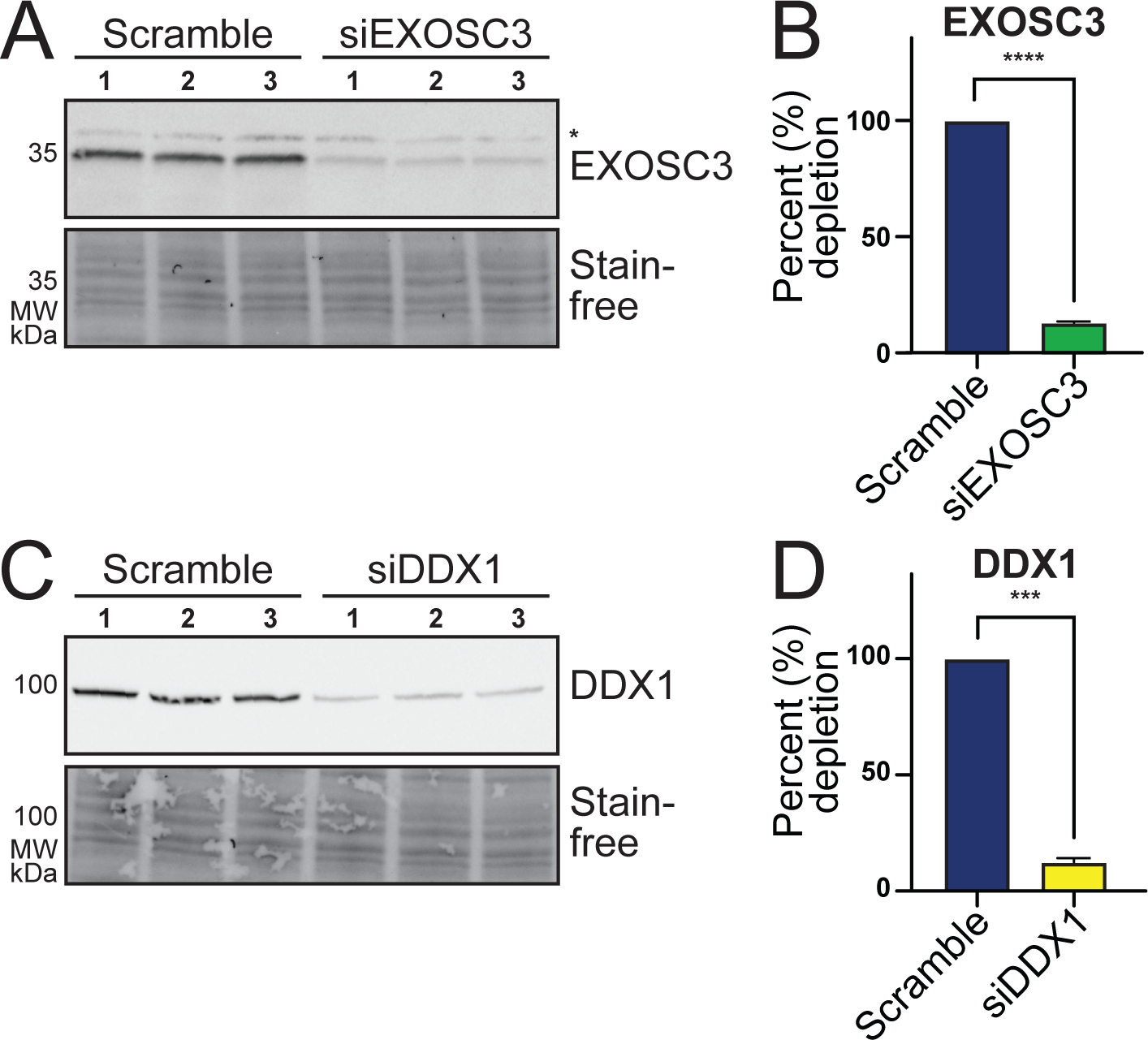
EXOSC3 and DDX1 are robustly depleted by siRNA-mediated knockdown in N2A cells. (A) N2A cells were transfected with Scramble control or EXOSC3 siRNA (siEXOSC3) in biological triplicate. The steady-state level of EXOSC3 was assessed by immunoblotting. The asterisk (*) indicates a nonspecific band. Stain-free blot serves as a loading control. (B) Quantification of immunoblot in (A) shows that EXOSC3 is depleted to 13.1%. The percent depletion of EXOSC3 was determined by quantifying the immunoblot in (A) relative to Scramble and averaging the values. This result is significant across three biological replicates. (C) N2A cells were transfected with Scramble or DDX1 siRNA (siDDX1) in biological triplicate. The steady-state level of DDX1 was assessed by immunoblotting. Stain-free blot serves as a loading control. (D) Quantification of immunoblot in (C) shows that DDX1 is depleted to 12.4%. The percent depletion of DDX1 was determined by quantifying the immunoblot in (C) relative to Scramble and averaging the values. This result is significant across three biological replicates. The statistical analyses for (C) and (D) were calculated using a student’s T-test. Asterisks (****) represent a p-value < 0.0001 and (***) represent a p-value < 0.001.

The RNA exosome has a well-defined role in rRNA processing and maturation (1,2,21,22,68). DEAD-box helicases such as DDX1 also play a critical role in RNA processing and genome stability (65,69). A previous study that analyzed DDX1 knockout mouse embryonic stem cells (mESCs) employed a pulse-chase experiment utilizing [^3^H]-uridine-labeled samples and showed an accumulation of precursor 28S rRNA and mature 18S rRNA, suggesting a role for DDX1 in rRNA processing (70). We performed a detailed analysis to explore rRNA processing in cells depleted of either EXOSC3 or DDX1. Using northern blotting to detect specific ribosomal RNA precursors, we examined which rRNA species are affected upon loss of either EXOSC3 or DDX1. **Figure 6A** depicts the steps of murine rRNA processing from the early precursor 47S to mature rRNA. Ribosomal RNA processing begins with the 47S precursor, which generates several downstream precursors, including 32S and 12S (71,72). The 47S precursor also produces the 18S, 5.8S, and 28S mature rRNAs, as well as the internal transcribed spacers 1 and 2 (ITS1 and ITS2), and the 5’ and 3’ external transcribed spacers (5’ETS and 3’ETS). To capture these precursors, we used probes specific to the ITS2 sequence. **Figure 6B** shows that the steady-state levels of the 12S precursor increases when EXOSC3 is depleted compared to the Scramble control, consistent with the most well-defined role of the RNA exosome in 3’ trimming to produce mature 5.8S (1,2,22,29,68). In contrast, when cells are depleted of DDX1, there is a decrease in the steady-state levels of the 12S precursor compared to the Scramble control. When using probes to detect 5.8S rRNA in the same samples, we detect a decrease in 5.8S level after depletion of EXOSC3, but no significant change when DDX1 is depleted. The 7SL signal recognition particle (SRP) transcript, which is not a target of the RNA exosome (73), is used as a loading control. The northern blot data are quantified for all analyses in **Figure 6C** and normalized to the loading control. The data from Figure 6B and Figure 6C are also summarized within Figure 6A, as indicated by the up- and down-arrows to denote statistically significant increases or decreases in these RNA species. Although both EXOSC3 and DDX1 clearly have an impact on rRNA processing or maturation, the roles in this process appear to be independent of each other.

**Figure 6:**
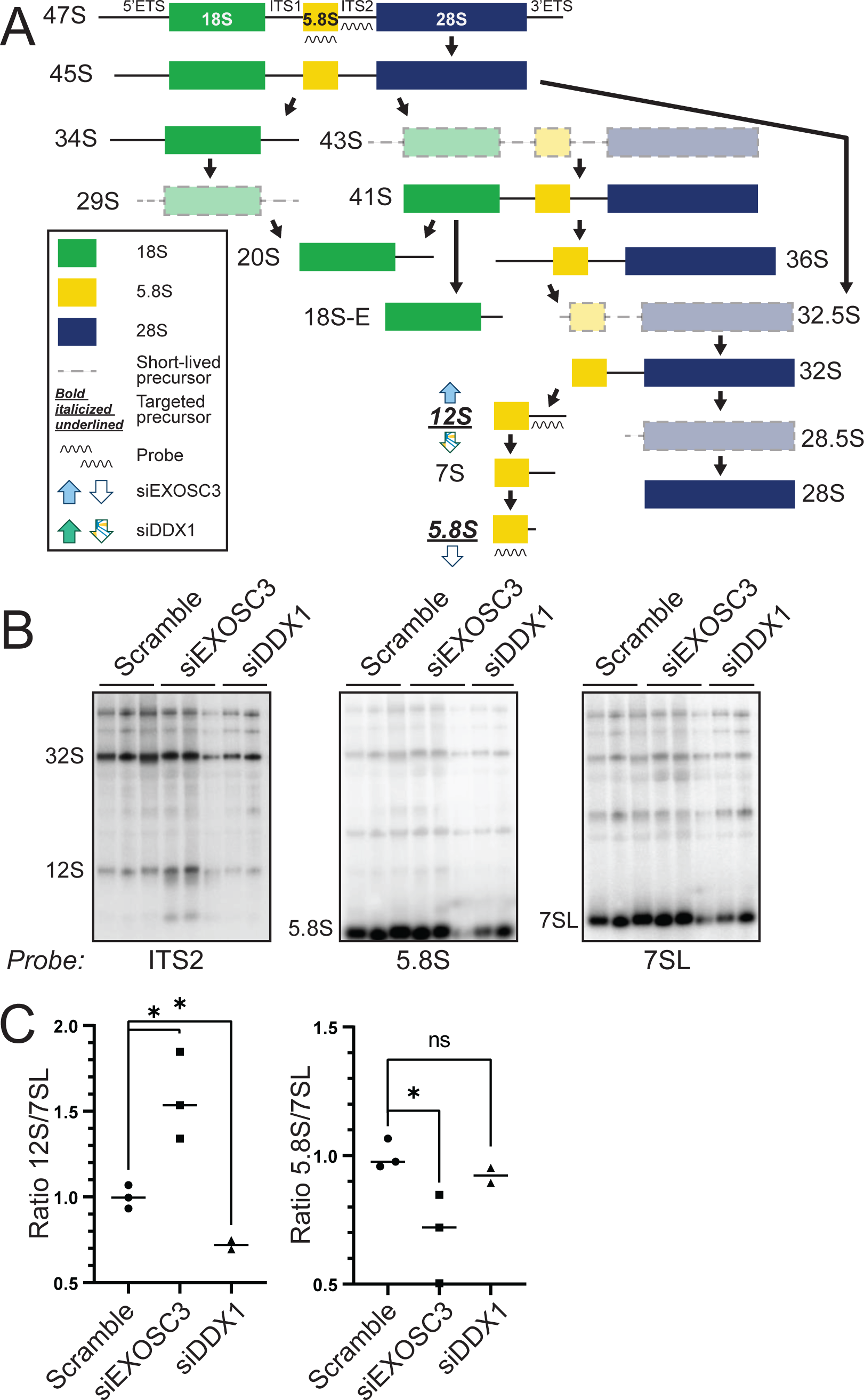
Depletion of EXOSC3 or DDX1 results in misprocessing of rRNA precursors. (A) A graphical schematic of murine rRNA processing, adapted from Henras *et al*., 2015 (71), that includes up- and down-arrows to summarize the results obtained in this study. (B) Northern blots of total RNA from N2A cells depleted of EXOSC3 or DDX1 using rRNA probes show that levels of 12S rRNA precursors and 5.8S rRNA are altered. EXOSC3 or DDX1 was depleted from cells by siRNA knockdown and total RNA was isolated for northern blotting. An ITS2 probe was used to detect 32S and 12S rRNA precursors. Additionally, we employed a probe specific for 5.8S rRNA. The 7SL transcript serves as a loading control. (C) Northern blots from Figure 6B were quantified relative to 7SL in biological triplicates. An asterisk (*) indicates a significant difference using a P-value cut-off of < 0.05.

### R-loops are globally reduced upon depletion of EXOSC3 or DDX1

Multiple studies have linked DDX1 to R-loops, which are three-strand nucleic acid structures comprised of an RNA-DNA hybrid and a single strand of DNA (58,69,74). DDX1 has been reported to co-precipitate with R-loops and promote R-loop formation by unwinding complex RNA (58,69,74). Studies have also linked the RNA exosome to R-loop regulation in murine B-cells (75). To understand the impact EXOSC3 and DDX1 may have on R-loops, we performed DNA/RNA-immunoprecipitation followed by high-throughput sequencing (DRIP-seq) in cells depleted of either EXOSC3 or DDX1. In cells depleted of EXOSC3, we identified 722 significantly increased R-loop regions and 935 decreased R-loop regions (**Figure 7A**, left, n = 3, FDR < 0.05). In cells depleted of DDX1, 638 increased R-loop regions and 1,058 decreased R-loop regions are identified (**Figure 7A**, right, n = 3, FDR < 0.05). We then compared the R-loop regions that showed statistically significant changes in cells depleted of EXOSC3 or DDX1. We found that 140 out of 722 increased R-loop regions (∼19%) in EXOSC3-depleted cells are also increased in DDX1-depleted cells (**Figure 7B**, left, Chi-squared test, p-value < 0.0005) and 425 out of 935 decreased R-loop regions (∼45%) in EXOSC3 knockdown cells are also decreased in DDX1 knockdown cells (**Figure 7B**, right, Chi-squared test, p-value < 0.0005). These results suggest that the two proteins may cooperate to alter a common set of R-loops.

**Figure 7:**
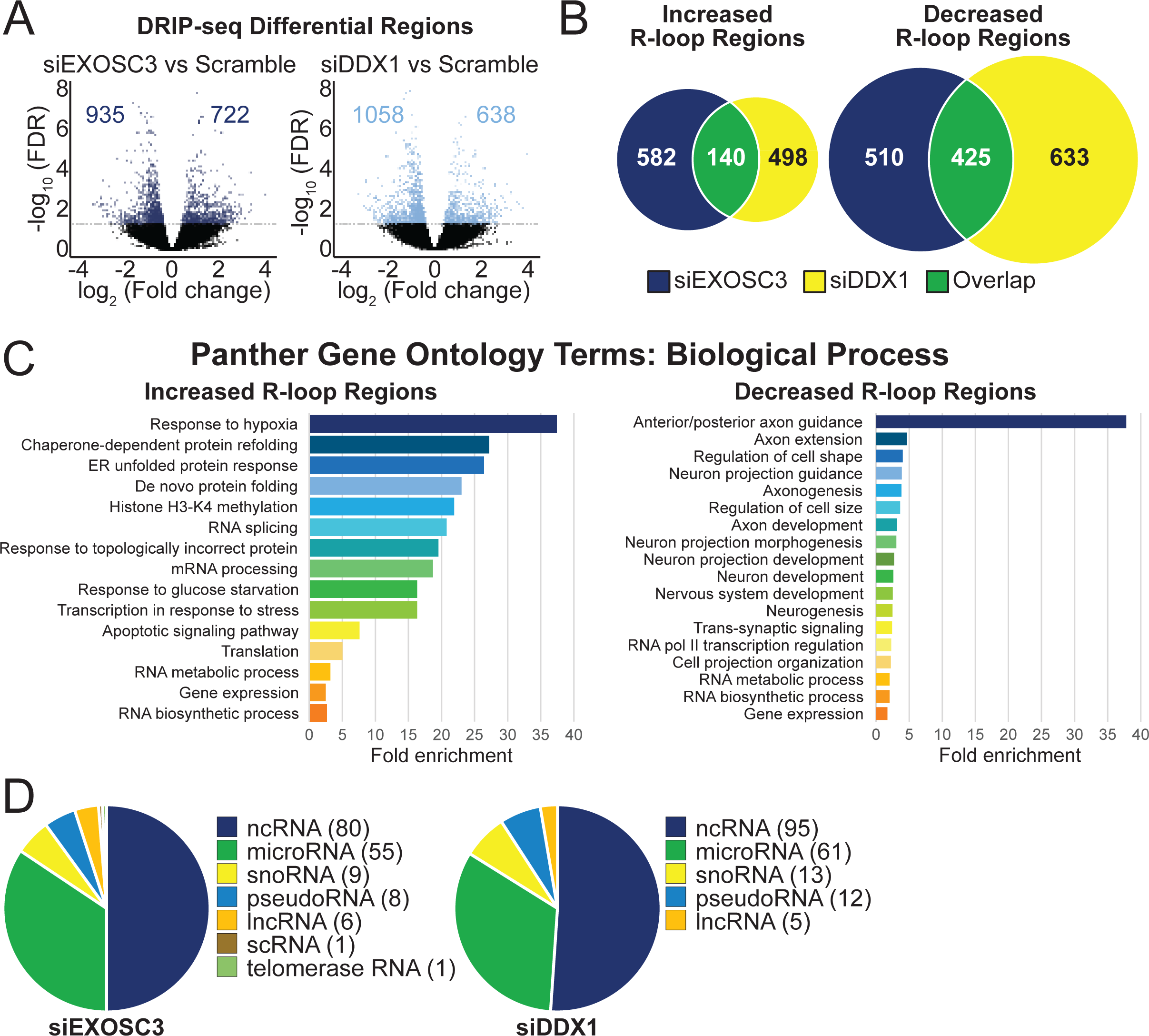
DRIP-seq reveals that depletion of EXOSC3 or DDX1 alters R-loop regions. DNA/RNA immunoprecipitation followed by sequencing (DRIP-seq) was performed on N2A cells depleted of EXOSC3 or DDX1 by siRNA. (A) The volcano plots show the number of R-loop regions that statistically increase and decrease in either EXOSC3 or DDX1 depletions compared to Scramble control. The plot is graphed using a log_2_ fold change across a -log_10_ false discovery rate (FDR). The R-loop regions that did not achieve the FDR cut-off of 0.05 are indicated in black and fall under the horizontal lines. The numbers on the left of 0 on the x axis indicate significantly decreased R-loop regions and the numbers on the right of 0 indicate significantly increased R-loop regions. (B) The statistically increased and decreased R-loop regions in cells siRNA depleted of EXOSC3 or DDX1 identified in the DRIP-seq dataset are compared using a Venn diagram. The dark blue circles indicate the number of increased (n = 722) or decreased (n = 935) R-loop regions upon siRNA-mediated EXOSC3 depletion. The yellow circles represent the number of increased (n = 638) or decreased (n = 1,058) R-loop regions that were affected upon siRNA-mediated DDX1 depletion. The overlap in green indicates the number of increased (n = 140) or decreased (n = 425) R-loop regions that siEXOSC3 or siDDX1 have in common. (C) The increased and decreased R-loop regions from both siEXOSC3 and siDDX1 samples were analyzed using Panther Gene Ontology Terms and categorized by biological process. These analyses are statistically significant with a cut-off at p-value < 0.05. (D) The classes of RNA that were affected upon depletion of either EXOSC3 or DDX1 are organized into a pie chart. Protein-coding genes were excluded. The numbers to the right of the class of RNA are the number of R-loop regions that correspond to that class.

EXOSC3- and DDX1-depleted cells share common R-loop regions that increased or decreased. We employed Panther Gene Ontology (GO) analysis on these R-loops (**Figure 7C**). The R-loop regions that are increased in both EXOSC3 and DDX1 knockdowns (n = 140) are enriched in the categories of protein folding, histone modifications, stress responses, and RNA metabolic processes when grouped by fold enrichment. The R-loop regions that are decreased in both EXOSC3 and DDX1 knockdown conditions (n = 425) are also enriched in the RNA metabolism category, but additionally in categories including axon guidance and neuronal development.

In **Figure 7D**, we grouped both increased and decreased R-loop regions by RNA transcript class present within the region corresponding to altered R-loops. Both EXOSC3 and DDX1 depletion affects R-loops within genes encoding different classes of RNA. We grouped all changed R-loop regions, both increased and decreased, after EXOSC3 depletion and found that the largest category of RNA significantly affected is protein-coding (n = 1,496). We then excluded protein-coding genes from the analysis to allow a clearer view of the non-coding transcripts and identified 80 ncRNAs, 55 microRNAs, 9 snoRNAs, 8 pseudoRNAs, 6 lncRNAs, 1 scRNA, and 1 telomerase RNA within the altered R-loop regions. An analysis following DDX1 depletion revealed similar results. We excluded 1,509 R-loop regions that mapped to protein-coding genes and identified 95 ncRNAs, 61 microRNAs, 13 snoRNAs, 12 pseudoRNAs, and 5 lncRNAs. Noncoding RNAs (ncRNAs) are those transcripts that do not currently have a more specific distinction.

In parallel with DRIP-seq, we employed RNA sequencing (RNA-seq) after ribosomal RNA depletion to identify transcripts altered by depletion of EXOSC3 or DDX1. The pipeline employed for this analysis focused on coding regions, so data presented represent changes in mRNA transcripts. We identified 1,757 significantly increased transcripts and 2,192 decreased transcripts in EXOSC3 knockdown cells (**Figure 8A**, left, n = 3, FDR < 0.05). In DDX1 knockdown cells, 734 increased transcripts and 968 decreased transcripts are identified (**Figure 8A**, right, n = 3, FDR < 0.05). We then compared the transcripts that showed statistically significant changes in EXOSC3- and DDX1-depleted cells. We found that 322 out of 1,757 increased mRNA transcripts (∼18%) in siEXOSC3 cells are also increased in siDDX1 cells (**Figure 8B**, left, Chi-squared test, p-value < 0.0005) and 599 out of 2,192 decreased mRNA transcripts (∼27%) in siEXOSC3 cells are also decreased in siDDX1 cells (**Figure 8B**, right, Chi-squared test, p-value < 0.0005), suggesting a coordination between EXOSC3 and DDX1 in commonly regulating a critical set of genes.

**Figure 8:**
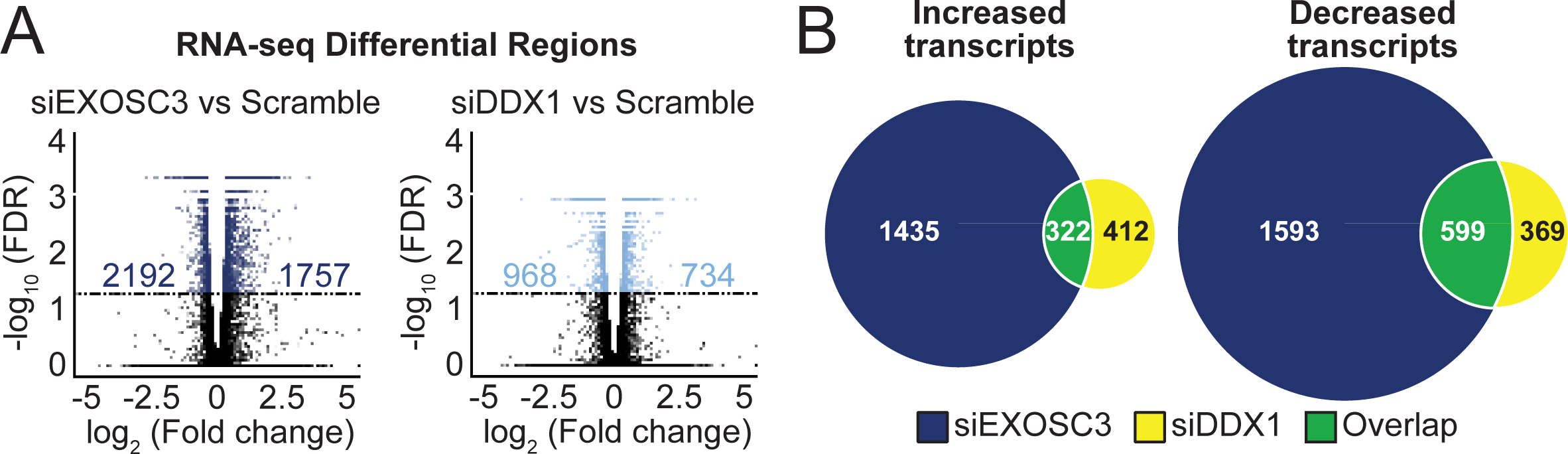
RNA-seq shows that depletion of EXOSC3 or DDX1 results in more shared decreased transcripts than shared increased transcripts. (A) RNA sequencing was performed on N2A cells siRNA depleted of EXOSC3 or DDX1. The volcano plots show the number of mRNA transcripts that increased and decreased in either siEXOSC3 or siDDX1 compared to Scramble control. The plot is graphed using a log_2_ fold change across a -log_10_ false discover rate (FDR). The differential regions that did not achieve the FDR cut-off of 0.05 are indicated in black and fall under the horizontal lines. The numbers on the left of 0 of the x axis indicate significantly decreased differential regions and the numbers on the right of 0 indicate significantly increased differential regions. (B) The increased and decreased transcripts are compared using a Venn diagram. The dark blue circles indicate the number of increased (n = 1,757) or decreased (n = 2,192) mRNA transcripts upon EXOSC3 depletion. The yellow circles represent the number of increased (n = 734) or decreased (n = 968) mRNA transcripts that are affected upon DDX1 depletion. The overlap represented in green indicates the number of increased (n = 322) or decreased (n = 599) mRNA transcripts that depletion of EXOSC3 or DDX1 have in common.

With both DRIP-seq and RNA-seq datasets in hand, we created a pipeline to compare results and filter the DRIP regions through the differential mRNA sequencing data to focus on the overlapping similarity between R-loops and changes in transcript level at those specific R-loop regions (**Figure 9A**). We applied the pipeline to generate a heatmap to illustrate the DRIP regions and mRNA transcripts that changed upon depletion of either EXOSC3 or DDX1 (**Figure 9B**). There are 466 R-loop regions that overlap in the RNA-sequencing data for both depletions. Of these overlapping R-loop regions, 103 are increased, and 363 regions are decreased. Many of the decreased regions mapped to genes that are involved in RNA metabolism, RNA regulation, translation processes, and neuronal development.

**Figure 9:**
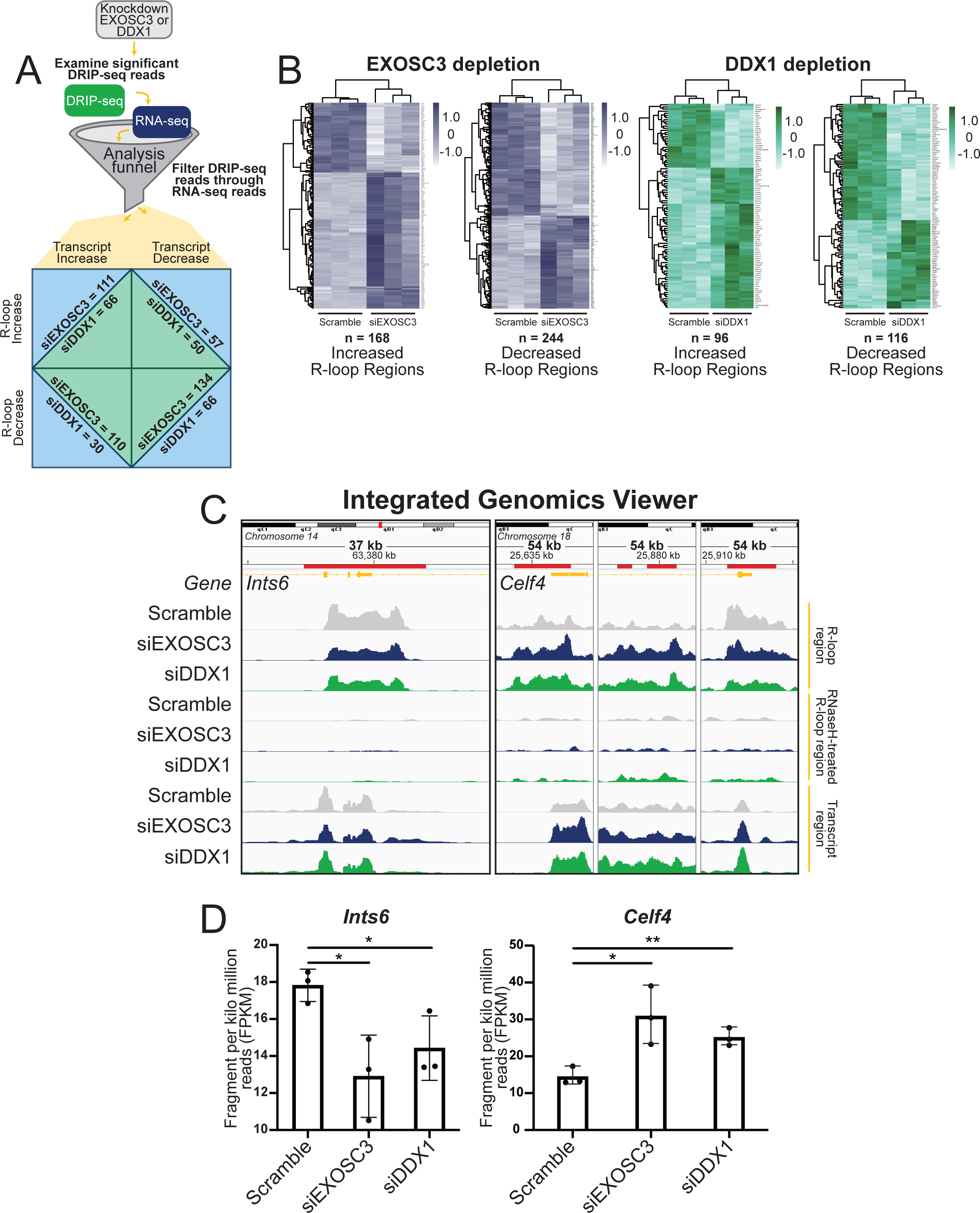
Filtering DRIP reads through RNA sequencing revealed genes that are simultaneously affected by depletions of EXOSC3 or DDX1. (A) A graphical representation of the pipeline used to focus on specific genes using the analysis funnel. N2A cells siRNA depleted of either EXOSC3 or DDX1 were subjected to both DRIP- and RNA-seq. We filtered results from the DRIP-seq through the RNA-seq reads, using only mRNA transcripts. We were then able to produce heatmaps and examine transcriptomic regions using an Integrated Genomics Viewer (IGV). (B) Using the pipeline described, we created heatmaps showing the landscape of increased and decreased R-loop regions upon depletion of either EXOSC3 (blue) or DDX1 (green). (C) The IGV images of *Ints6* and *Celf4*. The chromosome is displayed at the top of the window. The span lists the number of bases currently displayed. The tick marks indicate the chromosome locations. The red line marks the regions in which R-loops are significantly changed. The top track displays the *Mus musculus* reference genome (NCBI37/mm9) in orange. The following three tracks display the R-loop regions of interest corresponding to Scramble, siEXOSC3, and siDDX1, respectively. The middle three tracks display the RNase H-treated R-loop regions of interest corresponding to Scramble, siEXOSC3, and siDDX1, respectively. The last three tracks display the Transcript regions of interest corresponding to Scramble, siEXOSC3, and siDDX1, respectively. *Celf4* exhibited three changed regions in the genes and is displayed by separate panels. (D) Quantification of *Ints6* and *Celf4* transcripts in Scramble, siEXOSC3, and siDDX1 by fragment per kilo million reads (FPKM).

We employed the Integrative Genomics Viewer (IGV), which enables the visualization of these regions (**Figure 9C**). To illustrate the effect on specific loci, we focused on two genes that contained R-loop regions significantly changed within transcripts that are also significantly changed, and which fell under the biological processes aforementioned: *Ints6* and *Celf4*. The red marker above the gene (orange) in the IGVs of *Ints6* denotes the R-loop region, indicating a potential reduction in gene expression at these loci. In the three panels on the right, *Celf4* R-loop regions are marked with a red line above the gene (orange). The two left IGV panels corresponding to *Celf4* show increased R-loop and transcript regions. In the far right IGV panel corresponding to *Celf4*, the R-loop regions are decreased, and transcript regions are increased. The changes in R-loop regions at these loci indicate potential changes in gene regulation and expression upon the depletion of EXOSC3 or DDX1. All IGV panels show regions that have statistically significant changes compared to Scramble. Quantification of *Ints6* and *Celf4* transcripts show statistical significance (**Figure 9D**). Altogether, these data indicate that depletion of either EXOSC3 or DDX1 results in changes in R-loop regions that do not necessarily correspond to a similar change in transcript levels for that same gene.

## Discussion

In this study, we used a proteomic approach to identify RNA exosome-associated proteins in a neuronal cell line and identified an interaction between EXOSC3 and the putative RNA helicase DDX1. Although each protein is present in both the nucleus and the cytoplasm, the interaction was only detected in the nuclear fraction. To explore possible shared functions between EXOSC3 (RNA exosome) and DDX1 in which each of these proteins are implicated, we examined the EXOSC3-DDX1 interaction upon DNA damage, and effects on rRNA processing and on R-loop distribution upon depletion of EXOSC3 or DDX1. Our results suggest that EXOSC3 and DDX1 participate in shared regulation of R-loops and the transcripts produced within the genomic regions that form those R-loops. Taken together, these data define a potential mechanism by which an interaction between the RNA exosome complex and an RNA helicase could modulate R-loops.

The RNA exosome complex interacts with a number of proteins, termed cofactors, to confer specificity for different RNA targets (53). Many studies, including a number of elegant structures (5-8,10,49,76-78), show that the RNA exosome consists of a core set of subunits and cofactors that are present in one-to-one stoichiometry. The core RNA exosome then interacts with a number of different proteins to facilitate degradation or processing of many different RNA targets. These protein-protein binding events may be transient as the RNA exosome interacts with multiple cofactors at shared binding sites that have been revealed by biochemical and structural studies (7). Indeed, only a single peptide was identified in the mass spectrometry data in this study for the well-characterized RNA exosome cofactors, MTR4 (alternatively named SKIV2L2) and MPP6. These results provide evidence that the interactions between the RNA exosome and cofactors are likely to be dynamic and transient.

Beyond the previously characterized RNA exosome cofactors, this study identified several candidate RNA exosome-interacting proteins, which are located in different cellular compartments. For example, DDX1 is present in both the nucleus and the cytoplasm (64), while the Pumilio proteins, PUM1 and PUM2, are reported to control RNA stability exclusively in the cytoplasm (79). Thus, further studies could explore whether the interactions identified in whole cell lysate occur preferentially in one cellular compartment or another. Although DDX1 is readily detected in both the nucleus and the cytoplasm (**Figure 3B**) (64), robust interaction between EXOSC3 and DDX1 was only detected in the nuclear lysate. This compartment-specific interaction could mean that the RNA exosome and DDX1 interact in the context of chromatin. However, treatment with DNase I did not substantially decrease the EXOSC3-DDX1 association (**Figure S1C**). One possibility as to why the EXOSC3-DDX1 interaction is compartment-specific is that post-translational modifications (PTMs) could modulate binding. Studies in *S. pombe* have revealed PTMs in Dis3, Mtr4, Rrp40 (EXOSC3), Rrp43 (EXOSC8), and Rrp46 (EXOSC5), though only PTM mimetics in Dis3 and Mtr4 impacted RNA processing (80). No studies have explored whether PTMs regulate the function of DDX1. Such studies in the future could provide insight into how the RNA exosome complex dynamically interacts with so many different cofactors to target a large number of distinct RNAs.

Though we have identified and characterized the interaction between the RNA exosome cap subunit EXOSC3 and DDX1, we have not yet explored whether this interaction is direct or indirect. As EXOSC3 is a component of the RNA exosome complex, DDX1 may interact with EXOSC3 or with other RNA exosome subunits. Alternatively, as this interaction was identified through co-purification, the interaction could be indirect and mediated by RNA exosome cofactors or other proteins. While we detect all RNA exosome subunits co-purifying with EXOSC3, these experiments do not demonstrate that DDX1 interacts directly with any RNA exosome subunit beyond EXOSC3. The interaction between EXOSC3 and DDX1 increases significantly upon RNase treatment, particularly RNase A (**Figure 3C** and **Figure S2C**), a phenomenon suggested to occur for protein-protein interactions (65). One possible explanation for why the EXOSC3-DDX1 interaction increases upon digestion of RNA may be that RNA is bound between the two proteins, and potentially interfering or competing for the same binding site. In one conceivable model, DDX1 could unwind RNA for degradation/processing by the RNA exosome and the removal of RNA increases interaction with EXOSC3. A previous study reported a similar observation when examining the interaction between hnRNPK and DDX1 (65). Thus, future studies will be required to further characterize the interaction to test whether DDX1 could function as an RNA exosome cofactor like multiple other helicases.

A logical model to explain the interaction between the RNA exosome and DDX1 was that these factors could cooperate in response to DNA damage. The RNA exosome and DDX1 have both been implicated in double-strand break repair by homologous recombination (HR) and non-homologous end joining (NHEJ) (23,34,58,59). If these factors worked together to respond to DNA damage, we speculated that inducing DNA damage would increase this interaction. In contrast to this prediction, we found that double-strand breaks induced by treatment with the topoisomerase inhibitor camptothecin significantly reduced the interaction (**Figure 4B**). This result suggests that as DDX1 is recruited to sites of DNA damage (59,60), the interaction with the RNA exosome is lost. This finding led us to consider the possibility that the RNA exosome and DDX1 could share a function in cellular homeostasis in the absence of DNA damage. Alternatively, another model that cannot yet be eliminated is that the RNA exosome and DDX1 could cooperate to respond to specific types of DNA damage, which have not yet been tested.

In addition to DNA damage response, both the RNA exosome and DDX1 have been implicated in rRNA processing. The best-defined role for the RNA exosome is 3’ trimming to produce mature 5.8S RNA (1,2,22,68), while DDX1 has been implicated in the accumulation of a number of rRNA species (70). Analysis of rRNA processing in cells depleted of either EXOSC3 or DDX1 revealed that loss of these proteins alter some shared rRNA species, albeit in distinct manners (**Figure 6**). For example, the mouse precursor of mature 5.8S rRNA, 12S rRNA, accumulates in cells depleted of EXOSC3 with a concomitant decrease in mature 5.8S. In contrast, depletion of DDX1 led to a decrease in the level of the 12S rRNA precursor. While these results show that some of the same rRNAs are impacted by the loss of either EXOSC3 or DDX1, the impact is different.

Helicases belonging to the DEAD- and DExH-box families such as DDX1 and MTR4 play important roles in RNA processing beyond ribosomal RNA. R-loops are also a common target of these helicases. Though R-loops are necessary for cellular maintenance, these structures can pose a threat to the genome if they accumulate (35,81). DEAD/DExH-helicases are critical for resolving and regulating R-loops as they unwind the nucleic acid structures for subsequent degradation by ribonucleases. For example, DDX1 has been reported to unwind G-quadruplex structures that can stabilize R-loops during transcription (58). MTR4, a well-characterized nuclear cofactor, unwinds R-loops and degrades RNA in complex with the RNA exosome (82). Despite data suggesting that both DDX1 and MTR4 could help resolve R-loops, a number of differences between the helicases exist. For example, the SPRY protein binding domain in the N-terminus upstream of two helicase domains (**Figure 3A**) is unique to DDX1. The SPRY domain in DDX1 is inserted between a phosphate-binding P-loop motif and a single-strand DNA binding Ia motif, separating the motifs by 240 residues, instead of the usual 20-40 residues seen in other DEAD-box proteins (54). Examination of the SPRY domain of DDX1 compared with the protein-protein interface of MTR4 and the RNA exosome may reveal surface sites of interaction and would potentially explain why a SPRY domain lies upstream of two helicase domains within the DDX1 protein. Structures indicate that MTR4 interfaces with MPP6, which tethers the helicase to the RNA exosome cap (7,8). This interface could be a shared docking site for these helicases. Further studies on whether DDX1 and MTR4 may have some common functions would also shed light on the cellular roles of these proteins.

Many reports link either the RNA exosome or DDX1 to the modulation of R-loops (34,58,74,75,81). However, no reports to date directly compare the RNA exosome and DDX1 in R-loop modulation. One previous study analyzed a well-characterized R-loop region within the *BAMBI* gene by mass spectrometry and provided a detailed list of the proteins that co-immunoprecipitated with R-loops in this region (74). DDX1, EXOSC3, and several other subunits of the RNA exosome were detected in the association with this *BAMBI* R-loop. Consistent with this previous study, the DRIP-seq performed here identified R-loop regions in the *BAMBI* gene significantly altered in cells depleted of either EXOSC3 or DDX1 (**Figure S4**). These findings support a model where, at least for the *BAMBI* R-loop region, both EXOSC3 and DDX1 are co-located and may contribute to R-loop modulation.

We discovered several genes for which R-loop regions are significantly altered in the same manner by depletion of either EXOSC3 or DDX1. We coupled the genome-wide DRIP-seq analysis with RNA-seq analysis to focus on regions where changes in R-loops are associated with changes in transcript levels. This analysis revealed several common findings. While depletion of either EXOSC3 or DDX1 caused both some increases and some decreases in the number of R-loops detected, data showed larger numbers of decreased R-loop regions as compared to increased R-loop regions (**Figure 7A**). Considering these shared changes, GO analysis revealed that the genes located in regions that show increased R-loops map to genes implicated in a variety of cellular responses to stress, which could reflect a requirement for proper RNA exosome and/or DDX1 function to support normal cell physiology. Strikingly, the only GO term enriched more than 5-fold among regions with decreased R-loops is “Anterior/posterior axon guidance” (Fold enrichment > 35). This finding could suggest that these regions of the genome are less accessible and perhaps less actively transcribed when either EXOSC3 or DDX1 is depleted, suggesting a possible link to the neurological disorders that are caused by missense mutations in genes encoding structural subunits of the RNA exosome.

To define potential shared transcript targets of EXOSC3 and DDX1, we also performed RNA-seq analysis using a pipeline that distinguished protein-coding transcripts from other RNAs. This analysis revealed that more transcripts show a significant change (FDR < 0.05) in levels upon depletion of EXOSC3 (3,949 transcripts) as compared to DDX1 (1,702 transcripts). Notably, the percentage of transcripts increased (56% for siEXOSC3; 57% for siDDX1) and decreased (44% for siEXOSC3; 43% for siDDX1) are similar in both cases; in fact, decreased transcripts slightly outnumber increased transcripts for both depletions. Interestingly, the data reveal a large number of shared transcripts that are regulated, suggesting a shared role in modulating a subset of transcripts or a shared cellular response at the transcript level. To integrate the DRIP-seq and RNA-seq and provide insight into how changes in R-loops could correlate with altered gene expression within the region of the genome where the R-loop is located, we also narrowed the analysis to only consider regions where both R-loops and transcript levels were changed when either EXOSC3 or DDX1 was depleted with a focus on the shared changes. This analysis provided interesting sets of data to consider. One notable point is that the overall patterns of R-loop and transcript changes displayed in the heatmaps (**Figure 9B**) appear similar for both EXOSC3 and DDX1 depletions with regions of increased R-loops sharing more increased transcripts and conversely for regions of decreased R-loops. To delve into the data in more detail, we identified altered R-loops linked to RNA metabolism, DNA repair, and/or neurodevelopment. Two examples from this analysis are *Ints6* and *Celf4*. *Ints6* encodes for a protein that is a noncatalytic member of the Integrator complex, a protein complex involved in the processing of snRNAs, resolving lncRNAs, and metabolizing mRNAs (83,84). The Integrator complex, containing INTS6, is involved in transcription termination of lncRNAs that are synthesized at sites of double-strand breaks, also known as damage-induced long noncoding RNAs (dilncRNAs) (83,85). *Celf4* encodes for an RNA binding protein that regulates mRNA stability and binds 3’UTRs (86). *Celf4* has also been linked to neurodevelopmental disorders and expression is enriched in the central nervous system (87,88). These transcripts may be regulated by the RNA exosome and DDX1 in neurons and dysregulation could contribute to the neurological pathology that occurs in many exosomopathy patients. These two examples represent just a subset of the many altered genomic regions and transcripts that could contribute to cellular dysfunction.

The overall goal of this study was to identify RNA exosome interacting proteins present in a neuronal cell line with the underlying hypothesis that neuronal-specific interacting partners could be lost in exosomopathies, which show primarily neuronal pathologies despite the ubiquitous expression of the RNA exosome. While the identification of an interaction with an additional RNA helicase is exciting, DDX1, like the RNA exosome, is ubiquitously expressed. DDX1 has not been definitively linked to any monogenic disease, but there is a report within the undiagnosed disease network of an individual with a missense mutation (p.Thr280Arg), which would fall within the catalytic domain of DDX1. This individual is reported to suffer from seizures and developmental regression, which could suggest that DDX1, like the RNA exosome, is required for normal function in some regions of the brain. In summary, we identified and characterized a new interacting partner of the RNA exosome, and studies thus far suggest that DDX1 could interact with the RNA exosome to modulate R-loop accumulation at some loci and to regulate transcript levels of some shared transcripts. Taken together, this study suggests the interaction between the RNA exosome and RNA helicase DDX1 may be required for neuronal-specific gene regulation.

## Materials and Methods

### Cell culture and siRNA transfections

A mouse neuroblastoma cell line, Neuro2A (N2A) (ATCC) was maintained in a humidified incubator with 5% CO_2_ at 37°C in Dulbecco’s modified Eagle’s medium (DMEM) supplemented with 10% FBS, 100 units/mL penicillin G and 100 mg/mL streptomycin antibiotics (1%). Individual siRNAs were transfected into cultured cells using Lipofectamine® 2000 (Invitrogen) and Opti-MEM® (ThermoFisher) according to the manufacturer’s protocol in antibiotic-free media. Three 60 mm plates were seeded for each experimental condition. Each 60 mm plate was further seeded into 3 wells in 6-well plates. The 6-well plates were further processed for immunoblotting, DRIP-seq, or RNA-seq. Therefore, each biological replicate for all conditions underwent the same analysis. The siRNAs employed were: Scramble negative control (IDT; 51-01-14-04), *EXOSC3* siRNA 1 (IDT; mm.Ri.Exosc3.13.1), and *EXOSC3* siRNA 2 (IDT; mm.Ri.Exosc3.13.5).

### Protein purification and antibody production

The *Exosc3* open reading frame encoding the mouse EXOSC3 protein was cloned into pGEX-6P-2 plasmid (GE Healthcare Life Sciences (now Cytiva)) to create an N-terminally glutathione-S-transferase (GST) tagged EXOSC3 construct. Recombinant GST-EXOSC3 was expressed in *Escherichia coli* Rosetta 2 (DE3). The GST fusion protein was purified by affinity chromatography on glutathione-Sepharose (GE Healthcare Life Sciences (now Cytiva)) in 300 mM NaCl, 50 mM HEPES/NaOH (pH 7.5), 5% glycerol and 2 mM beta-mercapto-ethanol (βME). The fusion protein was eluted by addition of 30 mM reduced glutathione. The GST tag was removed using PreScission protease (GE Healthcare Life Sciences (now Cytiva)), and the EXOSC3 protein was further purified by collecting the flowthrough of a second affinity chromatography on glutathione-Sepharose resin. The untagged protein was further purified on a Superdex200 size exclusion chromatography column equilibrated in 150 mM NaCl, 30 mM HEPES/NaOH (pH 7.5), 5% glycerol. Expression and purification of both recombinant GST-EXOSC3 and untagged EXOSC3 were confirmed by SDS-PAGE followed by Coomassie staining. The purified untagged protein was used as an immunogen to raise rabbit polyclonal antibodies by Josman, LLC. Sera containing anti-EXOSC3 antibodies was collected 21 days after immunization and used directly for immunoblotting, immunoprecipitation, and immunofluorescence.

### Cell fractionation

N2A cells were collected, spun down, and gently resuspended in cold fractionation buffer (10 mM Tris-HCl pH 7.4, 10 mM NaCl, 0.5% v/v IGEPAL® CA-630 (Sigma)) supplemented with Pierce™ Protease Inhibitor Mini Tablet (ThermoFisher A32955) for 10 min on ice. Cell lysates were centrifuged at 1000xg for 5 minutes and the supernatant/cytoplasmic fraction was collected. The pellet was washed once with fractionation buffer and centrifuged at 1000xg for 5 minutes. The resulting pellet was collected as the nuclear fraction. Protein samples were resuspended in IP buffer (50 mM Tris-HCl pH 7.4, 100 mM NaCl, 0.5% IGEPAL® CA-630 (Sigma)) supplemented with Pierce™ Protease Inhibitor Mini Tablet (ThermoFisher A32955) and prepared for immunoblotting and immunoprecipitation.

### Immunoblotting

Protein lysates were boiled in reducing sample buffer and resolved on 4-20% Criterion TGX polyacrylamide gels (Bio-Rad), then transferred to 0.2 µm nitrocellulose membranes (Bio-Rad). Membranes were incubated for at least one hour in blocking buffer containing 5% non-fat dry milk in 0.5% Tris-buffered saline with Tween® (TBST). Membranes were then incubated for 1–2 hours at room temperature or overnight at 4°C in primary antibody diluted in blocking buffer. Primary antibodies were detected using species-specific horse radish peroxidase (HRP) conjugated secondary antibodies (Jackson ImmunoResearch) followed by incubation with enhanced chemiluminescence substrate (ECL) (Sigma) or SuperSignal™ West Femto Maximum Sensitivity Substrate (ThermoFisher). Chemiluminescence was detected by exposing blots using a ChemiDoc Imaging System (Bio-Rad). Immunoblots were quantified using ImageLab software compatible with the ChemiDoc Imaging System (Bio-Rad). Primary antibodies and dilutions as they appear: myc (mouse monoclonal; Cell Signaling 2276S; 1:1000), EXOSC9 (rabbit polyclonal; Bethyl A303-888A; 1:2000), EXOSC3 (rabbit polyclonal; custom made; 1:2000), HSP90 (Santa Cruz sc-13119; 1:400), B23 (mouse monoclonal; Santa Cruz sc-271737; 1:200), DDX1 (rabbit polyclonal; Bethyl A300-521A; 1:2000).

### Immunoprecipitation

*Mass spectrometry sample preparation:* N2A cells were transfected with pcDNA3 plasmids containing 2X myc-EXOSC3 or pcDNA3 empty vector (Addgene) using Lipofectamine^®^ 2000 (Thermofisher) and Opti-MEM^®^ (Thermofisher) as recommended by the manufacturer. Cell pellets were resuspended and lysed in 20 mM HEPES pH 7.2, 100 mM NaCl, and 0.2% Triton-X-100. Cell lysate was cleared with centrifugation for 10 minutes at 10,000 RPM at 4°C. Protein concentrations were determined using a standard Pierce™ bicinchoninic acid (BCA) assay (ThermoFisher, 23225). Total lysate (30 µg) was collected for input. For each sample, 20 µL Pierce™ Anti-c-Myc magnetic beads (Thermofisher #88842) were added to 600 µL-700 µL of lysate and incubated at room temperature for 1 hour while tumbling end over end. Beads were washed 3X and submitted for liquid chromatography and tandem mass spectrometry.

*Endogenous IP sample preparation:* Nuclear and cytoplasmic protein lysates were prepared as described previously in cell fractionation, followed by suspending in IP buffer (50 mM Tris-HCl pH 7.4, 100 mM NaCl, 0.5% IGEPAL® CA-630 (Sigma)) with one Pierce™ Protease Inhibitor Mini Tablet (ThermoFisher A32955). Lysates were passed through a 27-gauge syringe 5 times and sonicated for 5 pulses three times. Lysates were centrifuged at 14,000xg for 15 minutes at 4°C and protein concentrations were determined using a standard Pierce™ bicinchoninic acid (BCA) assay (ThermoFisher, 23225). Lysate (30 µg) was collected for input. Dynabeads™ Protein G magnetic beads (Invitrogen; 10003D) were suspended in 200-400 µL phosphate-buffered saline with Tween ® (PBST) and incubated with rotation for 30 minutes with nonspecific rabbit IgG isotype control (Invitrogen, 31235) or an equal volume of EXOSC3 antibody (50 µg/mL) at room temperature. Clarified lysates were added to washed bead-antibody complexes. If required, enzymes were added to final lysates and washed bead-anybody complexes: RNase A (0.2 µg/µL; Invitrogen 12091-039), RNase T1 (5 U; Worthington Biochemical LS01435), RNase H (8 U; Invitrogen 18021-014), DNase I (5 U; Invitrogen 18068015), Benzonase (1:10,000; Santa Cruz sc202391). The samples were incubated at 4°C overnight while tumbling end over end. After incubation, the beads were magnetized and washed three times with cold IP buffer. The protein complexes were eluted with reducing sample buffer (250 mM Tris-HCl pH 7.4, 500 mM dithiothreitol (DTT), 10% SDS, 0.5% bromophenol blue, and 50% glycerol) and prepared for immunoblotting.

### Liquid chromatography and tandem mass spectrometry (LC-MS/MS)

#### Sample preparation

Digestion was performed on beads using a digestion buffer containing 50 mM NH_4_HCO_3_. The beads were then treated with 1 mM dithiothreitol (DTT) at 25°C for 30 minutes, followed by addition of 5 mM iodoacetamide (IAA) at 25°C for 30 minutes in the dark. Lysyl endopeptidase (Wako) was added to the mixture at a 1:50 (w/w) enzyme to protein ratio and digestion proceeded at 25°C overnight. Samples were further digested overnight with 1:50 (w/w) trypsin (Promega) at 25°C. Resulting peptides were desalted with a Sep-Pak C18 column (Waters) and dried under vacuum.

#### LC-MS/MS analysis

Dried peptide samples were resuspended in 15 µL of loading buffer (0.1% formic acid, 0.03% trifluoroacetic acid, 1% acetonitrile) and 3 µL was loaded and separated on a self-packed C18 (1.9 µm Dr. Maisch, Germany) fused silica column (25 cm x 75 µM internal diameter (ID); New Objective, Woburn, MA) by a Dionex Ultimate 3000 RSLCNano and monitored on a Fusion mass spectrometer (ThermoFisher Scientific, San Jose, CA). Elution was performed over a 120-minute gradient at a rate of 250 nL/min with buffer B ranging from 3% to 35% (buffer A: 0.1% formic acid in water, buffer B: 0.1 % formic acid in acetonitrile). The mass spectrometer cycle was programmed to collect at the top speed for 3 second cycles. The mass spectrometry scans (300-1500 m/z range, 200,000 AGC, 50 ms maximum ion time) were collected at a resolution of 120,000 at m/z 200 in profile mode and the HCD MS/MS spectra (1.5 m/z isolation width, 0.5 m/z offset, 30% collision energy, 10,000 AGC target, 35 ms maximum ion time) were detected in the ion trap. Dynamic exclusion was set to exclude previous sequenced precursor ions for 20 seconds within a 10-ppm window. Precursor ions with +1 and +8 or higher charge states were excluded from sequencing.

#### Data analysis

Spectra were searched using Proteome Discoverer 2.1 against mouse Uniprot database (53,289 target sequences – downloaded April 2015). Searching parameters included fully tryptic restriction and a parent ion mass tolerance (± 20 ppm). Product ion tolerance was 0.6 Da. Methionine oxidation (+15.99492 Da), asparagine and glutamine deamidation (+0.98480), and protein N-terminal acetylation (+42.03670) were variable modifications (up to 3 allowed per peptide); cysteine was assigned a fixed carbamidomethyl modification (+57.021465 Da). Percolator was used to filter the peptide spectrum matches to a false discovery rate of 1%.

### Immunofluorescence

N2A cells were fixed on coverslips with 4% PFA (Electron Microscopy Sciences) for 20 minutes at room temperature and permeabilized with 0.5% Triton-X-100 (Sigma) for 10 minutes at room temperature. Coverslips were then blocked with 5% BSA for 60 minutes at room temperature and incubated overnight at 4°C with primary antibody γH2AX (rabbit monoclonal; Cell Signaling 9718T, 1:400). Texas red species-specific secondary antibodies (Jackson ImmunoResearch Laboratories Inc.) were used at a 1:200 dilution. Images were obtained using an Olympus BX60 inverted brightfield microscope (Olympus Life Sciences) with a 100X objective and recorded with a Moment CMOS camera (Teledyne Photometrics). Fluorescence intensity in green, blue, and red channels were measured using an automated program pipeline using Ocular v2.0 Advanced Scientific Camera Control.

### Total RNA preparation

Total RNA was isolated from N2A cells using the TRIzol reagent (Invitrogen) according to the manufacturer’s protocol. RNA quality was measured by 1% agarose gel electrophoresis (**Figure S3A**) and by High Sensitivity (HS) DNA Assay (Agilent) (**Figure S3B**).

### Northern blot

Total RNAs from three biological replicates were separated on a 1% agarose/formaldehyde gel and transferred by capillarity overnight in 20X saline sodium citrate (3 M NaCl, 300 mM sodium citrate pH 7) onto a nylon membrane (Cytiva). Membranes were probed as indicated. Bands were quantified using Image Lab software (Bio-Rad). Probe sequences were purchased from IDT and are listed in Supporting Information **Table S2**.

### DNA/RNA immunoprecipitation (DRIP) and sequencing

Genomic DNA was extracted from ∼4 million N2A cells and fragmented with restriction enzymes (BsrGI, EcoRI, HindIII, SspI, XbaI, 30 U/each, NEB) following a published protocol (89). Fragmented nucleic acids were recovered by phenol-chloroform extraction. Fragmented gDNA (4 μg) was digested with 20 U of RNase H (NEB #M0297S) for 6 hours to serve as a negative control. Fragmented gDNA or RNase H-digested DNA (4 μg) were immunoprecipitated with 8 µg of S9.6 antibody (Millipore #MABE1095) overnight at 4°C in DRIP binding buffer (10 mM sodium phosphate, 140 mM sodium chloride, 0.05% (v/v) Triton-X-100, pH 7.0). DNA-antibody complex was then incubated with 40 µL of Protein G beads (Invitrogen #10003D) for 2 hours at 4°C, and beads were washed three times with DRIP binding buffer. Antibody-captured DNA-RNA hybrids were then eluted in (50 mM Tris pH 8.0, 10 mM EDTA, 0.5% (v/v) SDS) and subjected to phenol-chloroform extraction. The immunoprecipitated DNA-RNA hybrids were quantified by Qubit (ThermoFisher Scientific #Q32854). DNA-RNA hybrids (10 ng) were treated with 5 U of RNase H for 1 hour and then were sonicated into small fragments (∼100-500 bp) using a Covaris Focused-Ultrasonicator Me220. Fragmented DNA was blunted, 5′ phosphorylated and 3’ A-tailed by NEBNext Ultra II End Prep Enzyme Mix following the manufacturer’s instruction (New England Biolabs #E7645L). An adaptor was ligated, and USER enzyme was used for U excision to yield adaptor-ligated double-strand DNA. The DNA was then PCR-amplified using barcoded PCR primers (New England Biolabs #E7335L, #E7500L). After AMPure XP bead purification (Beckman Coulter #A63881) and Qubit quantification, the libraries were sent to Admera Health for quality analysis and sequencing with Illumina with a read length configuration of 150 PE.

### Library preparation and high-throughput sequencing

For the rRNA-depleted RNA-seq library, sample quality was assessed by Bioanalyzer 2100 Eukaryote Total RNA Pico (Agilent Technologies, CA, USA) and samples were quantified by Qubit RNA HS assay (ThermoFisher #Q32852). Ribosomal RNA depletion was performed with Ribo-zero rRNA Removal Kit (Illumina Inc., San Diego, CA) followed by NEBNext® Ultra™ II Nondirectional RNA Library Prep Kit for Illumina® per manufacturer’s recommendation. Library concentration was measured by qPCR, and library quality was evaluated by Tapestation High Sensitivity D1000 ScreenTapes (Agilent Technologies, CA, USA). Equimolar pooling of libraries was performed based on qPCR values. Libraries were sequenced on a HiSeq with a read length configuration of 150 PE, targeting 80 million total reads per sample (40 million in each direction).

### Gene ontology (GO) enrichment

#### Mass spectrometry

Proteins exhibiting a log_2_ ratio greater than 0 peptide spectra matched (PSM) were analyzed using Panther Classification system (v17.0) and those exhibiting a ratio less than zero PSM were excluded. Of the 1141 proteins that met these criteria, 751 were organized into the functional classification system protein class. The pie chart was created in Microsoft Excel.

#### DRIP-seq

Fold enrichment of the modules was determined using Panther Classification System (v17.0) (DOI: 10.5281/zenodo.6799722). The total set of increased R-loop peaks (140) and decreased R-loop peaks (425) for both siDDX1 and siEXOSC3 datasets were set as the client text box input. A Fisher’s exact test was used to annotate the data set into biological processes. We used a false discovery rate (FDR) and P-value cut-off at 0.05 and focused on biological processes pertaining to RNA metabolism and neuron development. Fold enrichment was graphed using Microsoft Excel.

### Bioinformatic analysis

#### DRIP-seq

DRIP-seq reads were aligned to mouse genome sequence (mm9) by Bowtie2 version 2.4.4 with default parameter (90). Aligned reads in bam file were sorted by genomic coordinate by SAMtools (91). Peaks for each library were identified by MACS2 version 2.2.6 with input bam file (92). DRIP-seq peaks identified from Scramble and siEXOSC3/siDDX1 were merged and reads were re-counted in the merged peak regions. Differential analysis was conducted by DESeq2 based on the reads count matrix in the merged peak regions and significantly changed regions were defined by FDR<0.05 (93). The differential regions were annotated to genes by annotatePeaks.pl from HOMER version 4.11 (94).

#### RNA-seq

RNA-seq reads were aligned to mouse genome (mm9) by Tophat version 2.1.0 with default parameter (95). Aligned reads in bam file were sorted by genomic coordinate by SAMtools (91). Gene differential expression (DE) analysis was conducted by Cuffdiff version 2.2.1 (96). Significant DE genes were defined by FDR<0.05. Gene Ontology analysis were performed by PANTHER (97).

#### Integrative Genomics Viewer

Transcriptomic regions were viewed using the Integrative Genomics Viewer (IGV) from https://software.broadinstitute.org; v2.15.2 (98). Three biological replicates of the depletion conditions were merged into a single track. The *Mus musculus* genome (NCBI37/mm9) was used as a reference.

### Statistical analysis

Comparisons between experimental groups were made using an unpaired student’s T-test. All data are presented as means and standard error of the mean (SEM) (error bars) for at least three independent experiments, unless otherwise indicated. Asterisks (*) indicate statistical significance at p-value < 0.05.

### Data availability

The mass spectrometry proteomics data are available in **Table S1**.

### Supporting information

This article contains supporting information.

## Acknowledgements

The authors thank current and past members of the Corbett lab group, including Dr. Milo B. Fasken and Dr. Maria C. Sterrett, for support, instruction, and discussion. We thank Dr. Ambro van Hoof for supportive feedback and funding to support collaborative studies of the RNA exosome. We thank the Emory Proteomics Core for their support and guidance.

## Author contributions

JLD and AHC for conceptualization, JLD, SWL, and YH for data curation, JLD, SK, and YH for formal analysis, JLD, AHC, DY, HG, and BY for funding acquisition, JLD, SWL, YH, and SK for investigation, JLD, SWL, AHC, RHSJ for methodology, AHC for project administration, JLD, AHC, HG, DY, and BY for resources, BY and YH for software, AHC for supervision, JLD for validation, JLD and YH for visualization, JLD for writing original draft, JLD, RHSJ, AHC for writing review and editing.

## Funding and additional information

This research was funded by grants from the National Institute of General Medical Sciences to Ambro van Hoof and AHC (R01GM130147), HG (1R35GM138123), and BY (AG078937), grants from the National Cancer Institute to DY (R01CA178999, R01CA254403, U54 CA274513), and a Synergy II Award from Emory Woodruff Health Sciences Center to HG, AHC, and BY. JLD was supported by a Diversity Supplement to R01GM130147. The content is solely the responsibility of the authors and does not necessarily represent the official views of the National Institutes of Health.

## References

1. Mitchell, P., Petfalski, E., and Tollervey, D. (1996) The 3’ end of yeast 5.8S rRNA is generated by an exonuclease processing mechanism. Genes & development 10, 502–513

2. Mitchell, P., Petfalski, E., Shevchenko, A., Mann, M., and Tollervey, D. (1997) The exosome: a conserved eukaryotic RNA processing complex containing multiple 3’-->5’ exoribonucleases. Cell 91, 457–466

3. Allmang, C., Kufel, J., Chanfreau, G., Mitchell, P., Petfalski, E., and Tollervey, D. (1999) Functions of the exosome in rRNA, snoRNA and snRNA synthesis. EMBO J 18, 5399–5410

4. Zinder, J. C., and Lima, C. D. (2017) Targeting RNA for processing or destruction by the eukaryotic RNA exosome and its cofactors. Gene & Development 31

5. Makino, D. L., Baumgärtner, M., and Conti, E. (2013) Crystal structure of an RNA-bound 11-subunit eukaryotic exosome complex. Nature 495, 70

6. Gerlach, P., Schuller, J. M., Bonneau, F., Basquin, J., Reichelt, P., Falk, S., and Conti, E. (2018) Distinct and evolutionary conserved structural features of the human nuclear exosome complex. Elife 7

7. Weick, E. M., Puno, M. R., Januszyk, K., Zinder, J. C., DiMattia, M. A., and Lima, C. D. (2018) Helicase-Dependent RNA Decay Illuminated by a Cryo-EM Structure of a Human Nuclear RNA Exosome-MTR4 Complex. Cell 173, 1663-+

8. Falk, S., Bonneau, F., Ebert, J., Kogel, A., and Conti, E. (2017) Mpp6 Incorporation in the Nuclear Exosome Contributes to RNA Channeling through the Mtr4 Helicase. Cell Rep 20, 2279–2286

9. Tomecki, R., Kristiansen, M. S., Lykke-Andersen, S., Chlebowski, A., Larsen, K. M., Szczesny, R. J., Drazkowska, K., Pastula, A., Andersen, J. S., Stepien, P. P., Dziembowski, A., and Jensen, T. H. (2010) The human core exosome interacts with differentially localized processive RNases: hDIS3 and hDIS3L. Embo j 29, 2342–2357

10. Zinder, J. C., Wasmuth, E. V., and Lima, C. D. (2016) Nuclear RNA Exosome at 3.1 A Reveals Substrate Specificities, RNA Paths, and Allosteric Inhibition of Rrp44/Dis3. Mol Cell 64, 734–745

11. Bonneau, F., Basquin, J., Ebert, J., Lorentzen, E., and Conti, E. (2009) The yeast exosome functions as a macromolecular cage to channel RNA substrates for degradation. Cell 139, 547–559

12. Makino, D. L., Schuch, B., Stegmann, E., Baumgartner, M., Basquin, C., and Conti, E. (2015) RNA degradation paths in a 12-subunit nuclear exosome complex. Nature 524, 54–58

13. Kadowaki, T., Schneiter, R., Hitomi, M., and Tartakoff, A. M. (1995) Mutations in nucleolar proteins lead to nucleolar accumulation of polyA+ RNA in Saccharomyces cerevisiae. Mol Biol Cell 6, 1103–1110

14. Hou, D., Ruiz, M., and Andrulis, E. D. (2012) The ribonuclease Dis3 is an essential regulator of the developmental transcriptome. BMC Genomics 13, 359

15. Lim, S. J., Boyle, P. J., Chinen, M., Dale, R. K., and Lei, E. P. (2013) Genome-wide localization of exosome components to active promoters and chromatin insulators in Drosophila. Nucleic acids research 41, 2963–2980

16. Boczonadi, V., Muller, J. S., Pyle, A., Munkley, J., Dor, T., Quartararo, J., Ferrero, I., Karcagi, V., Giunta, M., Polvikoski, T., Birchall, D., Princzinger, A., Cinnamon, Y., Lutzkendorf, S., Piko, H., Reza, M., Florez, L., Santibanez-Koref, M., Griffin, H., Schuelke, M., Elpeleg, O., Kalaydjieva, L., Lochmuller, H., Elliott, D. J., Chinnery, P. F., Edvardson, S., and Horvath, R. (2014) EXOSC8 mutations alter mRNA metabolism and cause hypomyelination with spinal muscular atrophy and cerebellar hypoplasia. Nature communications 5, 4287

17. Slavotinek, A., Misceo, D., Htun, S., Mathisen, L., Frengen, E., Foreman, M., Hurtig, J. E., Enyenihi, L., Sterrett, M. C., and Leung, S. W. (2020) Biallelic variants in the RNA exosome gene EXOSC5 are associated with developmental delays, short stature, cerebellar hypoplasia and motor weakness. Human molecular genetics 29, 2218–2239

18. Morton, D. J., Jalloh, B., Kim, L., Kremsky, I., Nair, R. J., Nguyen, K. B., Rounds, J. C., Sterrett, M. C., Brown, B., and Le, T. (2020) A Drosophila model of Pontocerebellar Hypoplasia reveals a critical role for the RNA exosome in neurons. PLoS Genet. 16, e1008901

19. Uhlén, M., Fagerberg, L., Hallström, B. M., Lindskog, C., Oksvold, P., Mardinoglu, A., Sivertsson, Å., Kampf, C., Sjöstedt, E., Asplund, A., Olsson, I., Edlund, K., Lundberg, E., Navani, S., Szigyarto, C. A.-K., Odeberg, J., Djureinovic, D., Takanen, J. O., Hober, S., Alm, T., Edqvist, P.-H., Berling, H., Tegel, H., Mulder, J., Rockberg, J., Nilsson, P., Schwenk, J. M., Hamsten, M., von Feilitzen, K., Forsberg, M., Persson, L., Johansson, F., Zwahlen, M., von Heijne, G., Nielsen, J., and Pontén, F. (2015) Tissue-based map of the human proteome. Science 347, 1260419

20. Schaeffer, D., Tsanova, B., Barbas, A., Reis, F. P., Dastidar, E. G., Sanchez-Rotunno, M., Arraiano, C. M., and van Hoof, A. (2009) The exosome contains domains with specific endoribonuclease, exoribonuclease and cytoplasmic mRNA decay activities. Nat Struct Mol Biol 16, 56–62

21. Schilders, G., Raijmakers, R., Raats, J. M., and Pruijn, G. J. (2005) MPP6 is an exosome-associated RNA-binding protein involved in 5.8S rRNA maturation. Nucleic acids research 33, 6795–6804

22. Schuller, J. M., Falk, S., Fromm, L., Hurt, E., and Conti, E. (2018) Structure of the nuclear exosome captured on a maturing preribosome. Science 360, 219–222

23. Pefanis, E., Wang, J., Rothschild, G., Lim, J., Chao, J., Rabadan, R., Economides, A. N., and Basu, U. (2014) Noncoding RNA transcription targets AID to divergently transcribed loci in B cells. Nature 514, 389–393

24. Delan-Forino, C., Schneider, C., and Tollervey, D. (2017) Transcriptome-wide analysis of alternative routes for RNA substrates into the exosome complex. PLoS Genet. 13, 26

25. Schneider, C., Kudla, G., Wlotzka, W., Tuck, A., and Tollervey, D. (2012) Transcriptome-wide analysis of exosome targets. Mol Cell 48, 422–433

26. Houseley, J., LaCava, J., and Tollervey, D. (2006) RNA-quality control by the exosome. Nat Rev Mol Cell Biol 7, 529–539

27. Wyers, F., Rougemaille, M., Badis, G., Rousselle, J. C., Dufour, M. E., Boulay, J., Régnault, B., Devaux, F., Namane, A., Séraphin, B., Libri, D., and Jacquier, A. (2005) Cryptic pol II transcripts are degraded by a nuclear quality control pathway involving a new poly(A) polymerase. Cell 121, 725–737

28. Preker, P., Nielsen, J., Kammler, S., Lykke-Andersen, S., Christensen, M. S., Mapendano, C. K., Schierup, M. H., and Jensen, T. H. (2008) RNA exosome depletion reveals transcription upstream of active human promoters. Science 322, 1851–1854

29. Klauer, A. A., and van Hoof, A. (2012) Degradation of mRNAs that lack a stop codon: a decade of nonstop progress. Wiley Interdiscip Rev RNA 3, 649–660

30. Fan, J., Kuai, B., Wu, G. F., Wu, X. D., Chi, B. K., Wang, L. T., Wang, K., Shi, Z. B., Zhang, H., Chen, S., He, Z. S., Wang, S. Y., Zhou, Z. C., Li, G. H., and Cheng, H. (2017) Exosome cofactor hMTR4 competes with export adaptor ALYREF to ensure balanced nuclear RNA pools for degradation and export. Embo J. 36, 2870–2886

31. Lejeune, F., Li, X., and Maquat, L. E. (2003) Nonsense-mediated mRNA decay in mammalian cells involves decapping, deadenylating, and exonucleolytic activities. Mol Cell 12, 675–687

32. van Hoof, A., Staples, R. R., Baker, R. E., and Parker, R. (2000) Function of the ski4p (Csl4p) and Ski7p proteins in 3’-to-5’ degradation of mRNA. Mol Cell Biol 20, 8230–8243

33. Rigby, R. E., and Rehwinkel, J. (2015) RNA degradation in antiviral immunity and autoimmunity. Trends Immunol 36, 179–188

34. Marin-Vicente, C., Domingo-Prim, J., Eberle, A. B., and Visa, N. (2015) RRP6/EXOSC10 is required for the repair of DNA double-strand breaks by homologous recombination. Journal of cell science 128, 1097–1107

35. Richard, P., and Manley, J. L. (2017) R loops and links to human disease. Journal of molecular biology 429, 3168–3180

36. Richard, P., Feng, S., and Manley, J. L. (2013) A SUMO-dependent interaction between Senataxin and the exosome, disrupted in the neurodegenerative disease AOA2, targets the exosome to sites of transcription-induced DNA damage. Genes & development 27, 2227–2232

37. Fasken, M. B., Morton, D. J., Kuiper, E. G., Jones, S. K., Leung, S. W., and Corbett, A. H. (2020) The RNA Exosome and Human Disease. Methods in molecular biology (Clifton, N.J.) 2062, 3–33

38. de Amorim, J., Slavotinek, A., Fasken, M. B., Corbett, A. H., and Morton, D. J. (2020) Modeling Pathogenic Variants in the RNA Exosome. RNA Dis 7

39. Wan, J., Yourshaw, M., Mamsa, H., Rudnik-Schoneborn, S., Menezes, M. P., Hong, J. E., Leong, D. W., Senderek, J., Salman, M. S., Chitayat, D., Seeman, P., von Moers, A., Graul-Neumann, L., Kornberg, A. J., Castro-Gago, M., Sobrido, M. J., Sanefuji, M., Shieh, P. B., Salamon, N., Kim, R. C., Vinters, H. V., Chen, Z., Zerres, K., Ryan, M. M., Nelson, S. F., and Jen, J. C. (2012) Mutations in the RNA exosome component gene EXOSC3 cause pontocerebellar hypoplasia and spinal motor neuron degeneration. Nat Genet 44, 704–708

40. Namavar, Y., Barth, P. G., Poll-The, B. T., and Baas, F. (2011) Classification, diagnosis and potential mechanisms in pontocerebellar hypoplasia. Orphanet journal of rare diseases 6, 50

41. Rudnik-Schoneborn, S., Senderek, J., Jen, J. C., Houge, G., Seeman, P., Puchmajerova, A., Graul-Neumann, L., Seidel, U., Korinthenberg, R., Kirschner, J., Seeger, J., Ryan, M. M., Muntoni, F., Steinlin, M., Sztriha, L., Colomer, J., Hubner, C., Brockmann, K., Van Maldergem, L., Schiff, M., Holzinger, A., Barth, P., Reardon, W., Yourshaw, M., Nelson, S. F., Eggermann, T., and Zerres, K. (2013) Pontocerebellar hypoplasia type 1: clinical spectrum and relevance of EXOSC3 mutations. Neurology 80, 438–446

42. Ivanov, I., Atkinson, D., Litvinenko, I., Angelova, L., Andonova, S., Mumdjiev, H., Pacheva, I., Panova, M., Yordanova, R., Belovejdov, V., Petrova, A., Bosheva, M., Shmilev, T., Savov, A., and Jordanova, A. (2018) Pontocerebellar hypoplasia type 1 for the neuropediatrician: Genotype-phenotype correlations and diagnostic guidelines based on new cases and overview of the literature. European journal of paediatric neurology : EJPN : official journal of the European Paediatric Neurology Society 22, 674–681

43. Radvanska, E., Pos, Z., Zatkova, A., Hyblova, M., Bauer, F., Szemes, T., Kadasi, L., and Radvanszky, J. (2022) Molecularly confirmed pontocerebellar hypoplasia in a large family from Slovakia with four severely affected children. Bratisl Lek Listy 123, 568–572

44. Di Donato, N., Neuhann, T., Kahlert, A. K., Klink, B., Hackmann, K., Neuhann, I., Novotna, B., Schallner, J., Krause, C., Glass, I. A., Parnell, S. E., Benet-Pages, A., Nissen, A. M., Berger, W., Altmuller, J., Thiele, H., Weber, B. H., Schrock, E., Dobyns, W. B., Bier, A., and Rump, A. (2016) Mutations in EXOSC2 are associated with a novel syndrome characterised by retinitis pigmentosa, progressive hearing loss, premature ageing, short stature, mild intellectual disability and distinctive gestalt. J Med Genet 53, 419–425

45. Burns, D. T., Donkervoort, S., Muller, J. S., Knierim, E., Bharucha-Goebel, D., Faqeih, E. A., Bell, S. K., AlFaifi, A. Y., Monies, D., Millan, F., Retterer, K., Dyack, S., MacKay, S., Morales-Gonzalez, S., Giunta, M., Munro, B., Hudson, G., Scavina, M., Baker, L., Massini, T. C., Lek, M., Hu, Y., Ezzo, D., AlKuraya, F. S., Kang, P. B., Griffin, H., Foley, A. R., Schuelke, M., Horvath, R., and Bonnemann, C. G. (2018) Variants in EXOSC9 Disrupt the RNA Exosome and Result in Cerebellar Atrophy with Spinal Motor Neuronopathy. Am J Hum Genet 102, 858–873

46. Somashekar, P. H., Kaur, P., Stephen, J., Guleria, V. S., Kadavigere, R., Girisha, K. M., Bielas, S., Upadhyai, P., and Shukla, A. (2021) Bi-allelic missense variant, p. Ser35Leu in EXOSC1 is associated with pontocerebellar hypoplasia. Clinical Genetics 99, 594–600

47. Yang, X., Bayat, V., DiDonato, N., Zhao, Y., Zarnegar, B., Siprashvili, Z., Lopez-Pajares, V., Sun, T., Tao, S., Li, C., Rump, A., Khavari, P., and Lu, B. (2019) Genetic and genomic studies of pathogenic EXOSC2 mutations in the newly described disease SHRF implicate the autophagy pathway in disease pathogenesis. Human molecular genetics

48. Morton, D. J., Kuiper, E. G., Jones, S. K., Leung, S. W., Corbett, A. H., and Fasken, M. B. (2018) The RNA exosome and RNA exosome-linked disease. RNA 24, 127–142

49. Kowalinski, E., Kogel, A., Ebert, J., Reichelt, P., Stegmann, E., Habermann, B., and Conti, E. (2016) Structure of a Cytoplasmic 11-Subunit RNA Exosome Complex. Mol. Cell 63, 125–134

50. Halbach, F., Rode, M., and Conti, E. (2012) The crystal structure of S. cerevisiae Ski2, a DExH helicase associated with the cytoplasmic functions of the exosome. Rna 18, 124–134

51. Kögel, A., Keidel, A., Bonneau, F., Schäfer, I. B., and Conti, E. (2022) The human SKI complex regulates channeling of ribosome-bound RNA to the exosome via an intrinsic gatekeeping mechanism. Mol Cell 82, 756–769.e758

52. Zhang, E., Khanna, V., Dacheux, E., Namane, A., Doyen, A., Gomard, M., Turcotte, B., Jacquier, A., and Fromont-Racine, M. (2019) A specialised SKI complex assists the cytoplasmic RNA exosome in the absence of direct association with ribosomes. Embo Journal 38

53. Schmid, M., and Jensen, T. H. (2019) The Nuclear RNA Exosome and Its Cofactors. Adv Exp Med Biol 1203, 113–132

54. Schmid, S. R., and Linder, P. (1992) D-E-A-D protein family of putative RNA helicases. Mol Microbiol 6, 283–291

55. Linder, P., Lasko, P. F., Ashburner, M., Leroy, P., Nielsen, P. J., Nishi, K., Schnier, J., and Slonimski, P. P. (1989) Birth of the DEAD box. Nature 337, 121–122

56. Godbout, R., Hale, M., and Bisgrove, D. (1994) A human DEAD box protein with partial homology to heterogeneous nuclear ribonucleoprotein U. Gene 138, 243–245

57. Suzuki, T., Katada, E., Mizuoka, Y., Takagi, S., Kazuki, Y., Oshimura, M., Shindo, M., and Hara, T. (2021) A novel all-in-one conditional knockout system uncovered an essential role of DDX1 in ribosomal RNA processing. Nucleic acids research 49, e40

58. Ribeiro de Almeida, C., Dhir, S., Dhir, A., Moghaddam, A. E., Sattentau, Q., Meinhart, A., and Proudfoot, N. J. (2018) RNA Helicase DDX1 Converts RNA G-Quadruplex Structures into R-Loops to Promote IgH Class Switch Recombination. Mol Cell 70, 650–662.e658

59. Li, L., Germain, D. R., Poon, H. Y., Hildebrandt, M. R., Monckton, E. A., McDonald, D., Hendzel, M. J., and Godbout, R. (2016) DEAD Box 1 Facilitates Removal of RNA and Homologous Recombination at DNA Double-Strand Breaks. Mol Cell Biol 36, 2794–2810

60. Li, L., Monckton, E. A., and Godbout, R. (2008) A role for DEAD box 1 at DNA double-strand breaks. Mol Cell Biol 28, 6413–6425

61. Passinen, S., Valkila, J., Manninen, T., Syvala, H., and Ylikomi, T. (2001) The C-terminal half of Hsp90 is responsible for its cytoplasmic localization. European Journal of Biochemistry 268, 5337–5342

62. Ye, K. Q. (2005) Nucleophosmin/B23, a multifunctional protein that can regulate apoptosis. Cancer Biology & Therapy 4, 918–923

63. Perez-Gonzalez, A., Pazo, A., Navajas, R., Ciordia, S., Rodriguez-Frandsen, A., and Nieto, A. (2014) hCLE/C14orf166 associates with DDX1-HSPC117-FAM98B in a novel transcription-dependent shuttling RNA-transporting complex. PloS one 9, e90957

64. Godbout, R., Packer, M., and Bie, W. (1998) Overexpression of a DEAD box protein (DDX1) in neuroblastoma and retinoblastoma cell lines. Journal of Biological Chemistry 273, 21161–21168

65. Chen, H. C., Lin, W. C., Tsay, Y. G., Lee, S. C., and Chang, C. J. (2002) An RNA helicase, DDX1, interacting with poly(A) RNA and heterogeneous nuclear ribonucleoprotein K. J Biol Chem 277, 40403–40409

66. Ryan, A. J., Squires, S., Strutt, H. L., and Johnson, R. T. (1991) Camptothecin cytotoxicity in mammalian cells is associated with the induction of persistent double strand breaks in replicating DNA. Nucleic acids research 19, 3295–3300

67. Kuo, L. J., and Yang, L.-X. (2008) γ-H2AX-a novel biomarker for DNA double-strand breaks. In vivo 22, 305–309

68. Pirouz, M., Munafò, M., Ebrahimi, A. G., Choe, J., and Gregory, R. I. (2019) Exonuclease requirements for mammalian ribosomal RNA biogenesis and surveillance. Nature structural & molecular biology 26, 490–500

69. Cargill, M., Venkataraman, R., and Lee, S. (2021) DEAD-Box RNA Helicases and Genome Stability. Genes (Basel) 12

70. Suzuki, T., Katada, E., Mizuoka, Y., Takagi, S., Kazuki, Y., Oshimura, M., Shindo, M., and Hara, T. (2021) A novel all-in-one conditional knockout system uncovered an essential role of DDX1 in ribosomal RNA processing. Nucleic Acids Research 49, e40–e40

71. Henras, A. K., Plisson-Chastang, C., O’Donohue, M. F., Chakraborty, A., and Gleizes, P. E. (2015) An overview of pre-ribosomal RNA processing in eukaryotes. Wiley Interdiscip Rev RNA 6, 225–242

72. Burman, L. G., and Mauro, V. P. (2012) Analysis of rRNA processing and translation in mammalian cells using a synthetic 18S rRNA expression system. Nucleic Acids Res 40, 8085–8098

73. Leung, E., and Brown, J. D. (2010) Biogenesis of the signal recognition particle. Biochemical Society Transactions 38, 1093–1098

74. Wang, I. X., Grunseich, C., Fox, J., Burdick, J., Zhu, Z., Ravazian, N., Hafner, M., and Cheung, V. G. (2018) Human proteins that interact with RNA/DNA hybrids. Genome research 28, 1405–1414

75. Laffleur, B., Lim, J., Zhang, W., Chen, Y., Pefanis, E., Bizarro, J., Batista, C. R., Wu, L., Economides, A. N., and Wang, J. (2021) Noncoding RNA processing by DIS3 regulates chromosomal architecture and somatic hypermutation in B cells. Nature genetics 53, 230–242

76. Wasmuth, E. V., Zinder, J. C., Zattas, D., Das, M., and Lima, C. D. (2017) Structure and reconstitution of yeast Mpp6-nuclear exosome complexes reveals that Mpp6 stimulates RNA decay and recruits the Mtr4 helicase. Elife 6

77. Puno, M. R., and Lima, C. D. (2018) Structural basis for MTR4-ZCCHC8 interactions that stimulate the MTR4 helicase in the nuclear exosome-targeting complex. Proc. Natl. Acad. Sci. U. S. A. 115, E5506–E5515

78. Liu, Q., Greimann, J. C., and Lima, C. D. (2006) Reconstitution, activities, and structure of the eukaryotic RNA exosome. Cell 127, 1223–1237

79. Zhang, M., Chen, D., Xia, J., Han, W., Cui, X., Neuenkirchen, N., Hermes, G., Sestan, N., and Lin, H. (2017) Post-transcriptional regulation of mouse neurogenesis by Pumilio proteins. Genes & development 31, 1354–1369

80. Telekawa, C., Boisvert, F. M., and Bachand, F. (2018) Proteomic profiling and functional characterization of post-translational modifications of the fission yeast RNA exosome. Nucleic acids research 46, 11169–11183

81. Crossley, M. P., Bocek, M., and Cimprich, K. A. (2019) R-loops as cellular regulators and genomic threats. Molecular cell 73, 398–411

82. Nair, L., Chung, H., and Basu, U. (2020) Regulation of long non-coding RNAs and genome dynamics by the RNA surveillance machinery. Nature reviews Molecular cell biology 21, 123–136

83. Nojima, T., and Proudfoot, N. J. (2022) Mechanisms of lncRNA biogenesis as revealed by nascent transcriptomics. Nature Reviews Molecular Cell Biology 23, 389–406

84. Tatomer, D. C., Elrod, N. D., Liang, D., Xiao, M.-S., Jiang, J. Z., Jonathan, M., Huang, K.-L., Wagner, E. J., Cherry, S., and Wilusz, J. E. (2019) The Integrator complex cleaves nascent mRNAs to attenuate transcription. Genes & development 33, 1525–1538

85. Zhang, F., Ma, T., and Yu, X. (2013) A core hSSB1–INTS complex participates in the DNA damage response. Journal of cell science 126, 4850–4855

86. Wagnon, J. L., Briese, M., Sun, W., Mahaffey, C. L., Curk, T., Rot, G., Ule, J., and Frankel, W. N. (2012) CELF4 regulates translation and local abundance of a vast set of mRNAs, including genes associated with regulation of synaptic function. PLoS genetics 8, e1003067

87. Halgren, C., Bache, I., Bak, M., Myatt, M. W., Anderson, C. M., Brøndum-Nielsen, K., and Tommerup, N. (2012) Haploinsufficiency of CELF4 at 18q12. 2 is associated with developmental and behavioral disorders, seizures, eye manifestations, and obesity. European Journal of Human Genetics 20, 1315–1319

88. Yang, Y., Mahaffey, C. L., Bérubé, N., Maddatu, T. P., Cox, G. A., and Frankel, W. N. (2007) Complex seizure disorder caused by Brunol4 deficiency in mice. PLoS genetics 3, e124

89. Sanz, L. A., and Chédin, F. (2019) High-resolution, strand-specific R-loop mapping via S9.6-based DNA-RNA immunoprecipitation and high-throughput sequencing. Nat Protoc 14, 1734–1755

90. Langmead, B., and Salzberg, S. L. (2012) Fast gapped-read alignment with Bowtie 2. Nat Methods 9, 357–359

91. Li, H., Handsaker, B., Wysoker, A., Fennell, T., Ruan, J., Homer, N., Marth, G., Abecasis, G., and Durbin, R. (2009) The Sequence Alignment/Map format and SAMtools. Bioinformatics 25, 2078–2079

92. Zhang, Y., Liu, T., Meyer, C. A., Eeckhoute, J., Johnson, D. S., Bernstein, B. E., Nusbaum, C., Myers, R. M., Brown, M., Li, W., and Liu, X. S. (2008) Model-based analysis of ChIP-Seq (MACS). Genome Biol 9, R137

93. Love, M. I., Huber, W., and Anders, S. (2014) Moderated estimation of fold change and dispersion for RNA-seq data with DESeq2. Genome Biol 15, 550

94. Duttke, S. H., Chang, M. W., Heinz, S., and Benner, C. (2019) Identification and dynamic quantification of regulatory elements using total RNA. Genome Res 29, 1836–1846

95. Kim, D., Pertea, G., Trapnell, C., Pimentel, H., Kelley, R., and Salzberg, S. L. (2013) TopHat2: accurate alignment of transcriptomes in the presence of insertions, deletions and gene fusions. Genome Biol 14, R36

96. Trapnell, C., Roberts, A., Goff, L., Pertea, G., Kim, D., Kelley, D. R., Pimentel, H., Salzberg, S. L., Rinn, J. L., and Pachter, L. (2012) Differential gene and transcript expression analysis of RNA-seq experiments with TopHat and Cufflinks. Nat Protoc 7, 562–578

97. Mi, H., Muruganujan, A., Ebert, D., Huang, X., and Thomas, P. D. (2019) PANTHER version 14: more genomes, a new PANTHER GO-slim and improvements in enrichment analysis tools. Nucleic Acids Res 47, D419–d426

98. Robinson, J. T., Thorvaldsdóttir, H., Winckler, W., Guttman, M., Lander, E. S., Getz, G., and Mesirov, J. P. (2011) Integrative genomics viewer. Nat Biotechnol 29, 24–26

