## Supporting information for "The RNA helicase DDX1 associates with the nuclear RNA exosome and modulates R-loops"

**Northern blot probe sequences**

| | |
|---|---|
| ITS2-32 | ACCCACCGCAGCGGGTGACGCGATTGATCG |
| 5.8S-113 | GCAAGTGCGTTCGAAGTGTC |
| 7SL | CAAAACTCCCGTGCTGATCA |

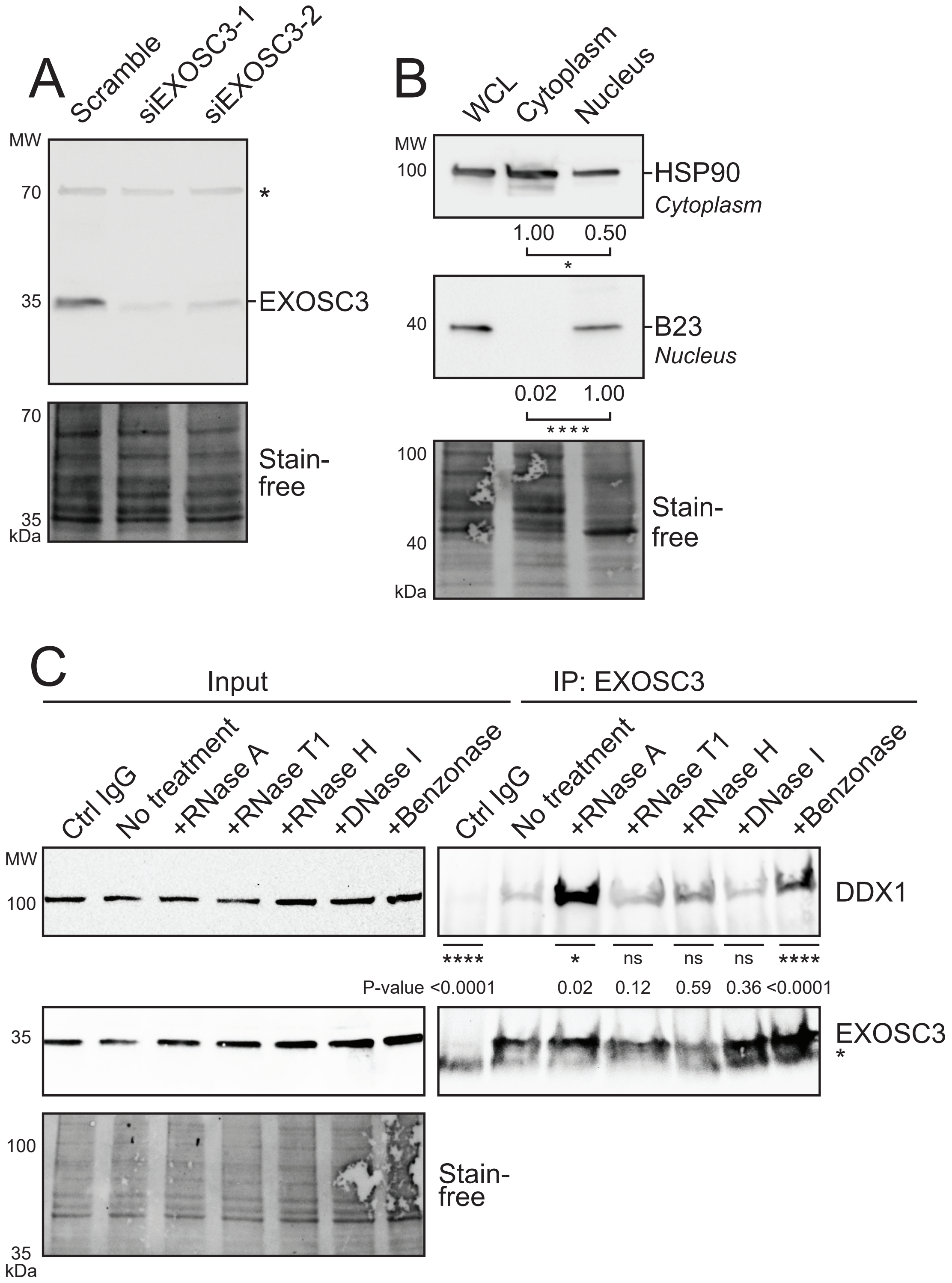
A
Scramble
siEXOSC3-1
siEXOSC3-2
MW
70
*
35
EXOSC3
70
Stain-free
35
kDa
B
WCL
Cytoplasm
Nucleus
MW
100
HSP90
Cytoplasm
1.00
0.50
*
40
B23
Nucleus
0.02
1.00
****
100
Stain-free
40
kDa
C
Input
IP: EXOSC3
Ctrl IgG
No treatment
+RNase A
+RNase T1
+RNase H
+DNase I
+Benzonase
Ctrl IgG
No treatment
+RNase A
+RNase T1
+RNase H
+DNase I
+Benzonase
MW
100
DDX1
****
*
ns
ns
ns
****
P-value <0.0001
0.02
0.12
0.59
0.36
<0.0001
35
EXOSC3
*
100
Stain-free
35
kDa

**Figure S1. EXOSC3 custom-made antibody is specific, cellular fractionation is sufficient, and the interaction between EXOSC3 and DDX1 is impacted by loss of RNA or DNA.**

(A) As described in Materials and Methods, polyclonal antibodies were created to detect EXOSC3. To validate the EXOSC3 antibody, N2A cells were transfected with Scramble or two independent EXOSC3 siRNAs (siEXOSC3-1 and siEXOSC3-2) and protein depletion was determined by immunoblot. The immunoblot was probed by the custom antibody. EXOSC3 antibody detects a band at the predicted size of EXOSC3, which is decreased with each EXOSC3 siRNA compared to Scramble. The Stain-free blot serves as a loading control for total protein. The asterisk (*) denotes a nonspecific band. (B) Cellular fractionation of N2A cells was performed as described in Materials and Methods. Equal amounts of protein from whole cell lysate (WCL), cytoplasmic, or nuclear fractions were analyzed with antibodies that detect proteins localized to either the cytoplasm (HSP90) or the nucleus (B23). The Stain-free blot serves as a loading control. The differences in protein profiles across the stain-free blot indicate the differences in protein populations between cytoplasmic and nuclear fractions. The bands are quantified relative to the nuclear (B23) or cytoplasmic (HSP90) marker in the respective cellular compartments. The values below the lanes correspond to the amount of protein quantified from the bands, normalized to the fraction observed; the cytoplasmic fraction was normalized to HSP90 (p-value = 0.0279) and the nuclear fraction was normalized to B23 (p-value < 0.0001). The quantification is averaged across biological triplicates and significance was calculated using student's T-test. (C) Immunoprecipitation of EXOSC3 from the nuclear fraction was performed with treatments of RNase A, RNase T1, RNase H, DNase I, and Benzonase. EXOSC3 antibody described previously is used in the No treatment, +RNase A, +RNase T1, +RNase H, +DNase I, and +Benzonase immunoprecipitation. Nonspecific rabbit IgG (Ctrl IgG) was used as a control. DDX1 and EXOSC3 were analyzed for the Input and bound fractions (IP: EXOSC3). The light chain IgG band is visible (asterisk) in the bound fractions probed with EXOSC3, just below the EXOSC3 band. Stain-free serves as a loading control for the Input. Bands in the bound fractions were quantified relative to the No treatment control. The values below the lanes correspond to the amount of protein quantified from the bands. This experiment was performed in duplicate (n = 2).

The asterisk (*) below the values indicate the P-value < 0.05.

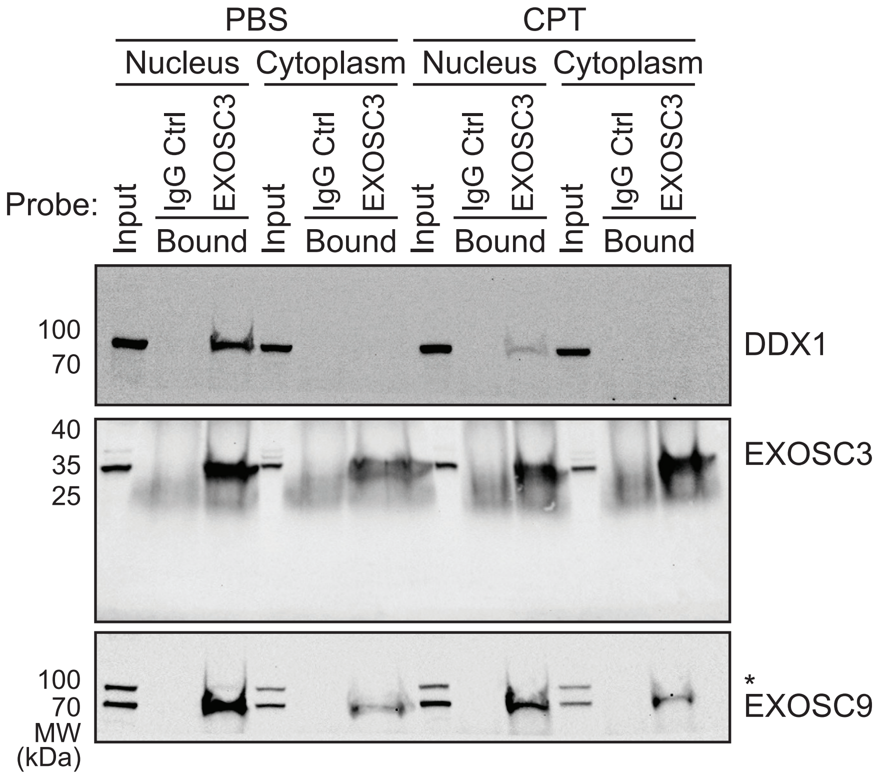
PBS
CPT
Nucleus
Cytoplasm
Nucleus
Cytoplasm
Probe:
Input
IgG Ctrl
EXOSC3
Bound
100
70
DDX1
40
35
25
EXOSC3
100
70
MW
(kDa)
*
EXOSC9

**Figure S2. The full immunoblot showing the interaction between EXOSC3 and DDX1 is DNA damage sensitive.**

(A) The immunoblot from Figure 4B is cropped from the immunoblot shown here. N2A cells were treated with PBS (control) or camptothecin (CPT) and prepared for immunoprecipitation as described in Materials and Methods. The Input and immunoprecipitated samples from nuclear and cytoplasmic fractions (Bound) treated with either PBS or CPT for both EXOSC3 and control IgG (Ctrl IgG) are shown. DDX1, EXOSC9, and EXOSC3 are detected. The immunoblot probed for DDX1 was also probed for EXOSC9. A band corresponding to DDX1 from the first probe is still visible and denoted with an asterisk (*) in the blot probed for EXOSC9.

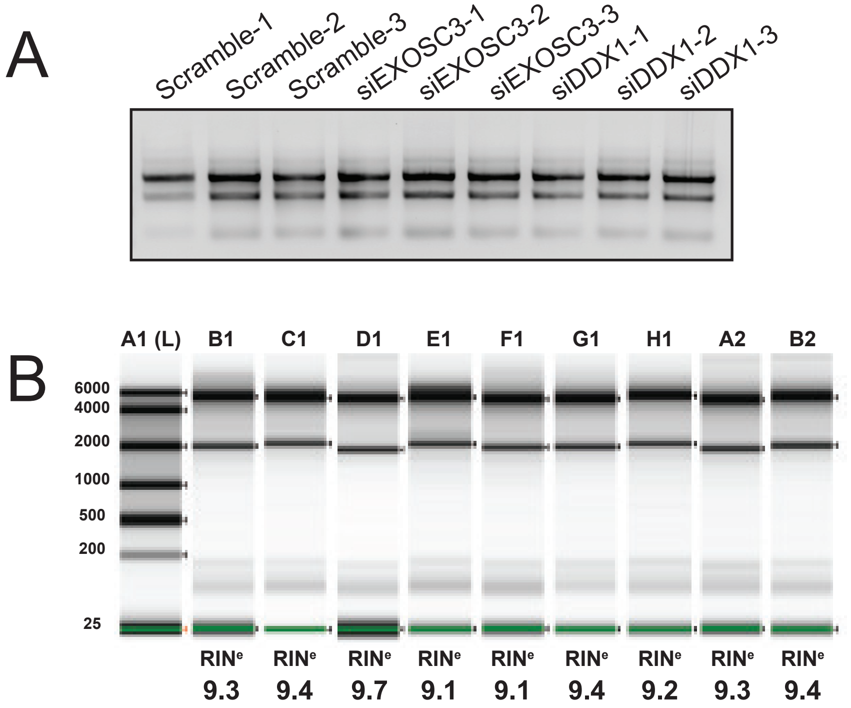


| Well | RINe | 28S/18S (Area) | Conc. [pg/µl] | Sample description |
|---|---|---|---|---|
| A1 (L) | - | - | 2140 | Ladder |
| B1 | 9.3 | 3.0 | 1880 | Scramble-1 |
| C1 | 9.4 | 3.5 | 13800 | Scramble-2 |
| D1 | 9.7 | 2.9 | 1050 | Scramble-3 |
| E1 | 9.1 | 3.8 | 4300 | siEXOSC3-1 |
| F1 | 9.1 | 3.2 | 2660 | siEXOSC3-2 |
| G1 | 9.4 | 3.1 | 4350 | siEXOSC3-3 |
| H1 | 9.2 | 2.7 | 3940 | siDDX1-1 |
| A2 | 9.3 | 3.0 | 2520 | siDDX1-2 |
| B2 | 9.4 | 3.0 | 2980 | siDDX1-3 |

**Figure S3. The quality of the purified RNA used for the northern blots, RNA-sequencing, and DRIP-sequencing assessed by 1% agarose gel and High Sensitivity ScreenTape assay.**

(A) RNA was extracted from N2A cells transfected with scramble siRNA, EXOSC3 siRNA, and DDX1 siRNA in biological triplicates. The quality of the RNA was assessed by a 1% agarose gel. The three bands visible correspond top to bottom to 28S, 18S, and 5.8S rRNA, respectively. (B) A High Sensitivity RNA ScreenTape assay was used for quantitation and further quality assessment. All RNA integrity numbers ($RIN^e$) for each sample are considered high quality.

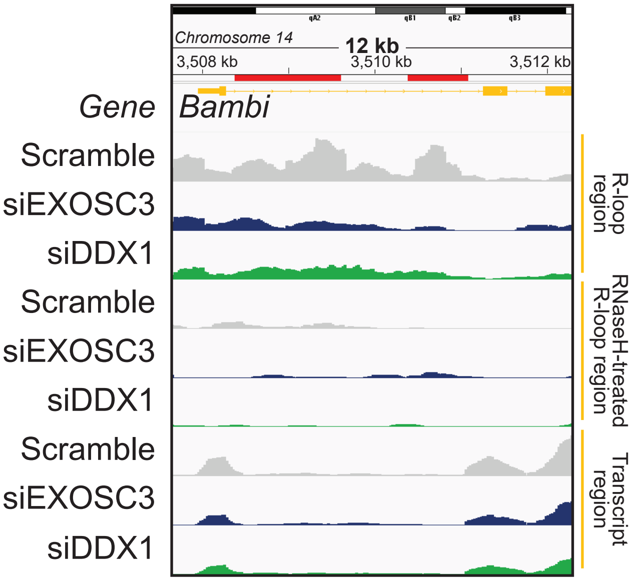

Chromosome 14
12 kb
3,508 kb
3,510 kb
3,512 kb
Gene
Bambi
Scramble
siEXOSC3
siDDX1
Scramble
siEXOSC3
siDDX1
Scramble
siEXOSC3
siDDX1
R-loop region
RNaseH-treated R-loop region
Transcript region

**Figure S4. IGV for R-loop regions in *Bambi* is elevated in cells depleted of *EXOSC3* or *DDX1.***

The Integrative Genome Viewer (IGV) image of *Bambi* is shown. The chromosome is displayed at the top of the window. The span lists the number of bases currently displayed. The tick marks indicate the chromosome locations. The red line marks the regions in which R-loops are significantly changed. The top track displays the Mus musculus reference genome (NCBI37/mm9) in orange. The following three tracks display the R-loop regions of interest corresponding to Scramble, siEXOSC3, and siDDX1, respectively. The middle three tracks display the RNase H-treated R-loop regions of interest corresponding to Scramble, siEXOSC3, and siDDX1, respectively. The last three tracks display the Transcript regions of interest corresponding to Scramble, siEXOSC3, and siDDX1, respectively.
